# Lysosomal membrane permeabilization enhances the anticancer effects of RNA Polymerase I transcription inhibitors

**DOI:** 10.1101/2025.03.03.641132

**Authors:** Lucille Ferret, Jonathan G Pol, Allan Sauvat, Gautier Stoll, Karla Alvarez-Valadez, Alexandra Muller, Julie Le Naour, Felix Peyre, Gerasimos Anagnostopoulos, Isabelle Martins, Maria Chiara Maiuri, Harald Wodrich, Lionel Guittat, Jean-Louis Mergny, Guido Kroemer, Mojgan Djavaheri-Mergny

**Affiliations:** Centre de Recherche des Cordeliers, Inserm UMRS 1138, Sorbonne Université, Université de Paris Cité, Équipe labellisée par la Ligue contre le Cancer, Institut Universitaire de France, 75006 Paris, France; Metabolomics and Cell Biology Platforms, Institut Gustave Roussy, 94805 Villejuif, France; Faculté de Médecine, Université de Paris Saclay, Paris, France; Department of Molecular Medicine and Medical Biotechnologies, University of Napoli Federico II, 80131 Naples, Italy; CNRS UMR 5234, Fundamental Microbiology and Pathogenicity, Université de Bordeaux, Bordeaux, France; Laboratoire d’Optique et Biosciences, École Polytechnique, CNRS UMR 7645-Inserm U 1182, Institut Polytechnique de Paris, 91120 Palaiseau, France; Université Sorbonne Paris Nord, UFR SMBH, Bobigny, France; Institut du Cancer Paris CARPEM, Department of Biology, Hôpital Européen Georges Pompidou, AP-HP, Paris, France

**Keywords:** autophagy, cancer, cell death, lysosome, resistance to therapy, transcription, TFEB

## Abstract

Lysosomes are known to contribute to the development of drug resistance through a variety of mechanisms that include the sequestration of drugs within their compartments and the activation of adaptive stress pathways. Although targeting POL I (RNA polymerase I) exhibits anticancer effects, little attention has been paid to the contribution of lysosomes to the efficacy and resistance of RNA POL I inhibitors. In this study, we investigated this aspect in the context of two potent POL I inhibitors, CX-3543 (Quarfloxin) and CX-5461 (Pidnarulex). Unexpectedly, CX-3543 was discovered to be sequestered in the lysosomal compartment. This resulted in the permeabilization of lysosomal membranes (LMP) and the subsequent activation of cellular stress adaptation pathways, including the transcription factor (TFEB) and autophagy. Disruption of TFEB or autophagy increased cell sensitivity to CX-3543, highlighting the cytoprotective role of these processes against cell death induced by this compound. Moreover, targeting lysosomal membranes using chloroquine derivatives or blue light excitation induced substantial LMP, resulting in the liberation of CX-3543 from lysosomes. This effect amplified both the inhibition of DNA-to-RNA transcription and cell death induced by CX-3543. Similar effects were observed when chloroquine derivatives were combined with CX-5461. Furthermore, combining CX-3543 with the chloroquine derivative DC661 reduced the growth of fibrosarcoma established in immunocompetent mice more efficiently than either agent alone. Altogether, our results uncover an unanticipated lysosome-related mechanism that contributes to the resistance of cancer cells to POL I transcription inhibitors, as well as a strategy to combat this resistance.

## Introduction

Lysosomes are essential organelles that regulate both catabolic and anabolic processes [1]. They serve as internal sensors of nutrient abundance and cellular stresses, activating various gene expression programs to modulate cell growth, cell death, and proliferation [2,3]. Major changes in lysosomes occur during malignant transformation and tumor progression [4,5]. Cancer-associated alterations in these organelles may allow to selectively target malignant cells, which are particularly dependent on lysosomal catabolic pathways such as endocytosis and macroautophagy/autophagy [6–8].

Accordingly, some anticancer therapies eliminate malignant cells by lysosome-dependent mechanisms. These include agents that provoke lysosomal membrane permeabilization (LMP), alter lysosomal pH, and disrupt the activity of lysosomal enzymes. These effects may ultimately cause cell death depending on the intensity of lysosome damage, the activation of adaptive stress responses, and the cell type [9]. In fact, the integrity of lysosomal membranes in cancer cells is often compromised, rendering them more vulnerable to lysosomal membrane damage than healthy cells [5,10]. Conversely, chemoresistance of cancer cells can be caused by sequestration of drugs within lysosomes, withholding them from their molecular targets [11–13]. This drug resistance can be overcome by treating cells with agents that raise the pH of lysosomes or cause lysosomal membrane permeabilization [9].

Upon LMP induction, cells can avoid cell death by activating adaptive processes to repair, recycle, and replace damaged lysosomes [14]. Suppression of these adaptive responses sensitizes cancer cells to LMP inducers. The Endosomal Sorting Complex Required for Transport (ESCRT) machinery plays a major role in lysosomal membrane repair. Unrepaired lysosomes can be degraded by selective autophagy, also called lysophagy. Of note, lysosomal membrane damage also results in the translocation of transcription factor EB (TFEB) to the nucleus, where TFEB then transactivates genes involved in autophagy and lysosomal biogenesis [15–17]. We previously demonstrated the unexpected capacity of DNA ligands that target G-quadruplex (G4) structures to induce autophagy and lysosomal biogenesis, as well as the possibility to sensitize cancer cells to such G4-ligands by lysosomotropic drugs [18–20]. G-quadruplexes are unusual nucleic acid structures composed of guanine-rich sequences. They have been the focus of extensive research in the cancer field due to their potential to control gene expression, DNA replication, RNA transcription and translation, and telomere functions [21]. CX-3543 and CX-5461 are prototypical G4-ligands that potently inhibit the transcriptional activity of RNA Polymerase I (POL I), thereby killing cancer cells [22]. Both agents exert promising anticancer effects in preclinical studies and have reached clinical trials [22–24]. However, until now, little attention has been given to the potential mechanisms involved in resistance to CX-3543 and CX-5461 and how such a resistance mechanism might be overcome.

Here, we took advantage of the fluorescence emission of CX-3543 to follow its subcellular distribution. Motivated by its unexpected lysosomal localization, we investigated the effects of CX-3543 on lysosomal integrity and function. We determined how enhancing lysosomal membrane permeabilization can potentiate its anticancer effects and carried out similar studies with CX-5461. Our findings uncovered a new mechanism through which CX compounds exert their cellular effects, as well as a novel combinatorial therapeutic approach to enhance their antineoplastic effectiveness.

## Results

### CX-3543 and CX-5461 trigger lysosomal membrane permeabilization

CX-3543 was first developed as a G4-ligand that inhibits RNA polymerase I [22]. Fluorescence microscopy analyses of human osteosarcoma U2OS cells treated with CX-3543 revealed that, upon blue light excitation, CX-3543 emits green fluorescence visible in discrete cytoplasmic puncta (**Figure 1A**). These puncta colocalized with Lysotracker red (a specific dye for acidic organelles such as lysosomes) but not with Mitotracker red (a mitochondrial dye), indicating that CX-3543 preferentially accumulates in the lysosomal compartment (**Figure 1B and C**).

**Figure 1.**
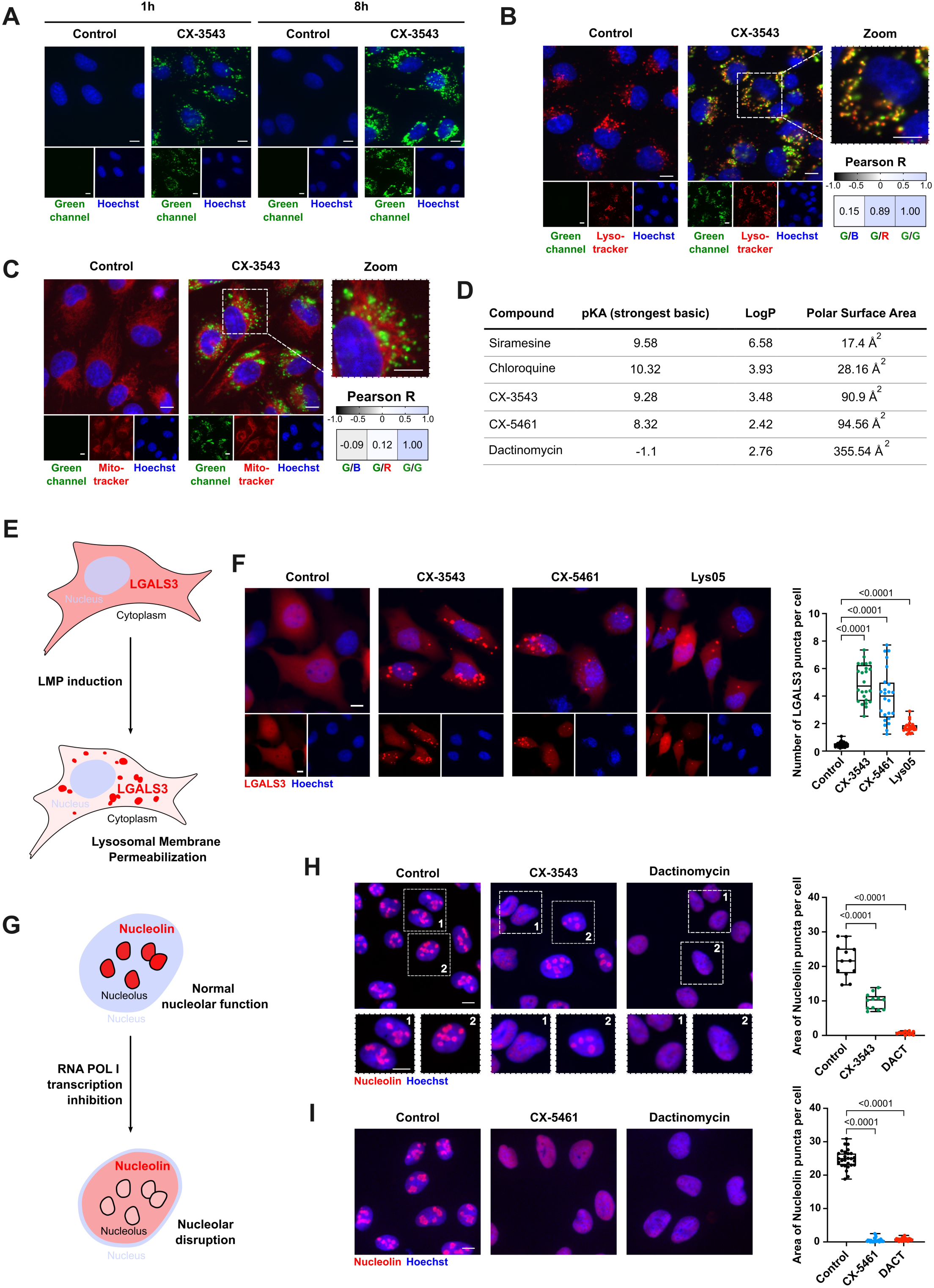
CX-3543 and CX-5461 promote lysosomal membrane permeabilization. **A)** To visualize the intrinsic fluorescence of CX-3543, human osteosarcoma U2OS cells were treated with CX-3543 (3 µM) during either 1 h or 8 h. The green fluorescence was then acquired by high-content microscopy (HCM) using a 40X objective. Hoechst 33342 was used to stain the nuclei. **B-C)** Human osteosarcoma U2OS cells were treated with 3 µM CX-3543 for 16 h followed by staining with specific subcellular markers for HCM analysis using a 40X objective. Lysosomes were visualized using Lysotracker™ Red DND dye (**B**), while mitochondria were stained by the MitoTracker™ Red dye (**C**). Representative confocal fluorescence images show CX-3543 fluorescence (green), Lysotracker (red) (**B**) or MitoTracker (red) (**C**), and nuclei (blue). The colocalization of CX-3543 with Lysotracker or MitoTracker was quantified using the Pearson Correlation Coefficient (R). **D)** Prediction of lysosomotropic molecular characteristics of CX-3543 and CX-5461 thanks to DrugBank online data. **E)** Schematic representation showing the shift of LGALS3 from a diffuse to a punctuate pattern following induction of lysosomal membrane permeabilization. **F)** U2OS cells stably expressing LGALS3-mCherry were treated with either CX-3543 (3 µM), or CX-5461 (5 µM) for 6 h. Lys05 (10 µM, 6 h) served as an inducer of LMP. HCM analysis was performed to evaluate the number of LGALS3 puncta values per cell with a minimum of 4,800 cells analyzed per condition. **G)** The diagram provides a schematic representation of the disruption of the nucleolus caused by the inhibition of Polymerase I transcription. Its highlights the relocation of the nucleolus marker, Nucleolin, from the nucleolus to the nucleoplasm. **H-I)** U2OS cells were treated with 3 µM CX-3543 or 5 µM CX-5461 for 16 h, then immunostained for detection of Nucleolin. Dactinomycin (1 µM, 2 h) served as a positive control. HCM analysis was performed to evaluate the Nucleolin area fluorescence intensity in the nucleolus per cell, with a minimum of 2,400 cells (**I**) or 1,700 cells (**J**) analyzed per condition. **Data analysis**: Statistical significance was calculated by means of a Pearson Correlation Coefficient R test (Figure 1B and C) or a Robust Linear Mixed model (Figure 1F-J). Scale bars: 10 μm.

The presence of CX-3543 within lysosomes is reminiscent of the behavior of lysosomotropic agents that accumulate in lysosomes and impair lysosomal membrane integrity and function [25]. To address this possibility, we conducted an *in silico* study to identify whether CX-3543 and CX-5461 possess physicochemical characteristics compatible with lysosomotropism, including a marked lipophilicity (assessed *via* the partition coefficient between water and n-octanol, logP) and a basic pKa value. Molecules with a logP > 2 and a pKa ranging from 6.5 to 11 tend to accumulate in lysosomes [25,26]. Using molecular descriptors retrieved from the Drugbank online database, both CX-3543 and CX-5461 were predicted to be lysosomotropic, similar to known lysosomotropic agents such as chloroquine and siramesine. In contrast, dactinomycin, another POL I transcription inhibitor, did not share these properties (**Figure 1D**).

High doses of lysosomotropic agents induce LMP [9,27]. We next investigated whether CX-3543 and CX-5461 trigger lysosome damage by means of a specific biosensor, LGALS3 (galectin 3). The LGALS3 protein is usually diffusely distributed in the cytosol, but is recruited to lysosomes undergoing LMP thanks to its affinity for glycans contained in the lumen of these organelles [28] (**Figure 1E)**. In the U2OS cell line stably expressing LGALS3-mCherry, exposure to CX-3543 or CX-5461 resulted in the appearance of red fluorescent puncta indicative of the translocation of LGALS3 to damaged lysosomes (**Figure 1F**). This LMP induction was further confirmed by immunofluorescent detection of endogenous LGALS3, which redistributed to puncta in U2OS and HeLa cells (Figure S1A and B).

Next, we explored the effect of CX-3543 on nucleolar integrity in U2OS cells immunostained for Nucleolin, a marker that is usually confined to nucleoli but spreads to the rest of the nucleoplasm following POL I inhibition (**Figure 1G**), as shown for dactinomycin [29–31]. In response to CX-3543 and even more to CX-5461, a subset of the cells displayed a dactinomycin-like nucleolar disruption phenotype (**Figures 1H and 1I**) suggesting partial nucleolar disruption under these treatments. Additionally, treatment with CX-3543 partially inhibited transcription as demonstrated by the 5-ethynyl uridine (5-EU) incorporation assay (Figure S1C). These findings not only align with previously reported POL I inhibitory effect of these compounds but also importantly reveal their lysosomotropic properties and ability to induce lysosomal membrane permeabilization (LMP).

### CX-3543 induces cell death through a caspase-independent mechanism

Upon LMP induction, cells can undergo cell death through caspase-dependent or independent mechanisms [9,10] (**Figure 2A**). CX-3543 triggered a dose-dependent loss of cell viability, with IC50 values of 4.9 µM in U2OS cells, 2.25 µM in HeLa cells and 1.6 µM in Jurkat cells (**Figure 2B**, Figure S2A-S2C). To further characterize the type of cell death, we used a pan-caspase inhibitor (Q-VD-OPh) and ferrostatin-1, which inhibit apoptosis and ferroptosis, respectively. As shown in **Figure 2C**, ferrostatin-1 did not prevent cell death induced by CX-3543, while it reduced cell death induced by the ferroptosis inducer erastin. Moreover, Q-VD-OPh failed to interfere with CX-3543-induced cell death, although it prevented cell death induced by the apoptosis inducer etoposide. Accordingly, the treatment with CX-3543 did not induce the cleavage of Caspase-3 whereas etoposide did (**Figure 2D**). Since lysosome-dependent cell death (LDCD) is characterized by the release of cathepsins, we investigated the potential effect of CX-3543 on LDCD effect by using cathepsins inhibitors, E64D and Pepstatin A. As shown in **Figure 2E**, the combination of E64D and Pepstatin A failed to inhibit CX-3543 triggered cell death. However, this combination effectively reduced cell death induced by LLOMe, a well-established inducer of lysosome-dependent cell death [32]. This finding demonstrates that cell death induced by CX-3543 occurs through a mechanism independent of LDCD.

**Figure 2.**
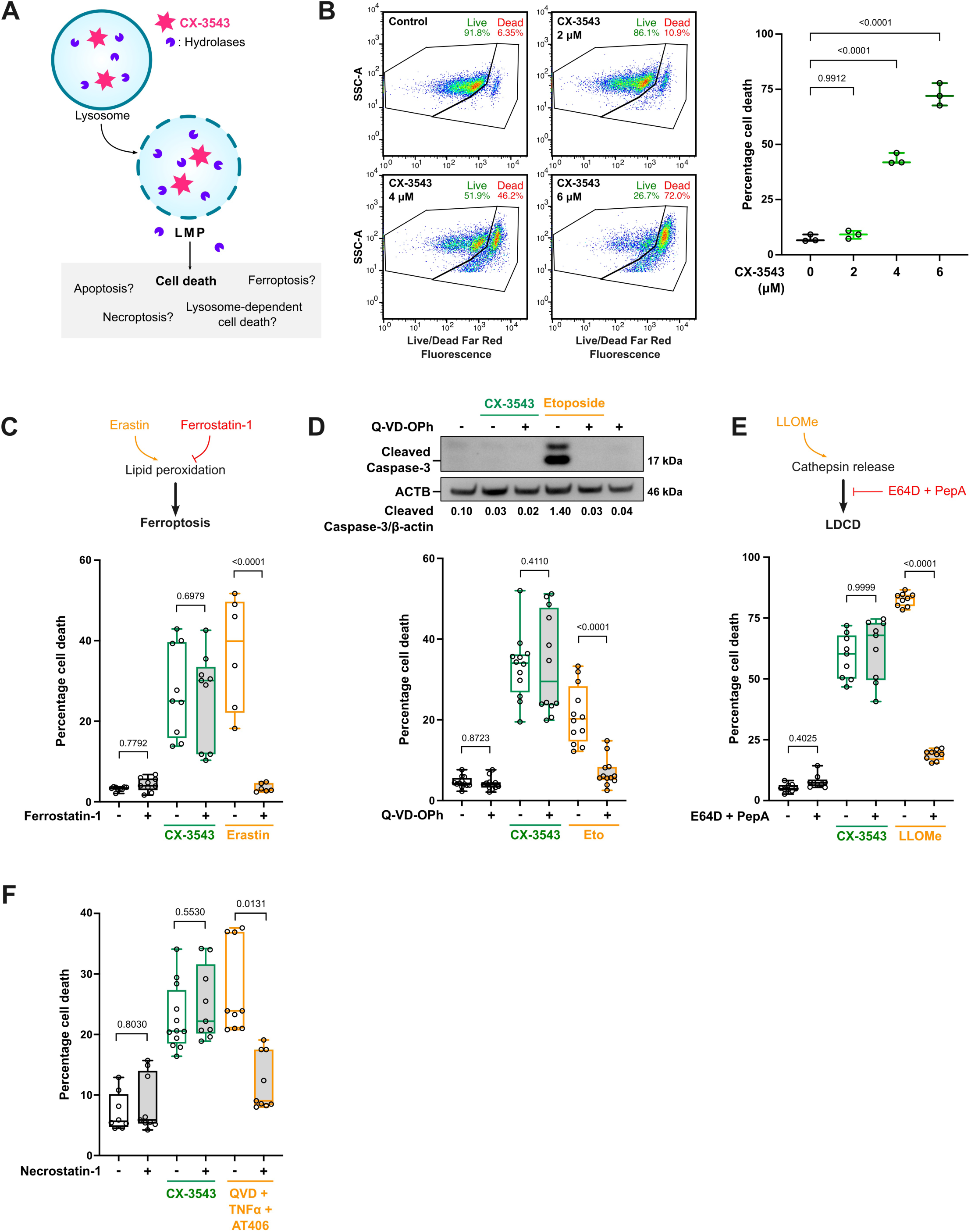
CX-3543 triggers cell death through a caspase-independent mechanism. **A)** A schematic representation depicting various types of cell death that can be triggered upon induction of lysosomal membrane permeabilization. **B)** U2OS cells were treated with indicated concentrations of CX-3543 for 24 h and then subjected to cell death detection by flow cytometry using a Live/Dead Far Red assay. **C) *Upper panel:*** a simplified presentation of ferroptosis regulation. ***Lower panel***: U2OS cells exposed to the ferroptosis inhibitor, ferrostatin-1 (5 µM), for 2 h prior to treatment with either 10 µM erastin (serving as a positive control for ferroptosis) or CX-3543 (3 µM) for 24 h. Cell death was then evaluated using a Live/Dead Far Red assay (N=3). **D)** U2OS cells were pre-treated with the pan-caspase inhibitor, Q-VD-OPh (40 µM), for 2 h before introducing CX-3543 (3 µM) or etoposide (100 µM) (serving as an inducer of apoptosis) for 24 h. ***Upper panel:*** Analysis of cleaved forms of Caspase-3 by immunoblot assay. ***Lower panel***: Cell death was evaluated using a Live/Dead Far Red assay (N=4). **E) *Upper panel:*** a simplified presentation of Lysosomal dependent cell death (LDCD) regulation. ***Lower panel***: U2OS cells were exposed to the cathepsins inhibitors, E64D (Cathepsin B and L) and Pepstatin A (Cathepsin D and E), for 2 h prior to treatment with either 2 mM LLOMe (serving as a positive control for LDCD induction) or CX-3543 (3 µM) for 24 h. Cell death was then assessed by a Live/Dead Far Red assay (N=3). **F)** To induce necroptosis as a positive control, Jurkat cells were pre-treated for 1 hour with a combination of the pan-caspase inhibitor Q-VD-OPh (10 µM), the SMAC mimetic AT406 (10 µM), and the necroptosis inhibitor necrostatin-1 (20 µM), followed by treatment with TNFα (50 ng/ml) for 24 hours. In parallel, Jurkat cells were exposed to necrostatin-1 for 1h prior treatment with CX-3543 (1 µM) for 24 h. Cell death was then assessed by a Live/Dead Far Red assay. **Data analysis**: Statistical significance was calculated by means of a one-coefficient robust linear model (**2B**) or a two-coefficients robust linear model (**2C-F**).

We next investigated the ability of CX-3543 to induce necroptosis using necrostatin-1, a specific inhibitor of this cell death pathway. The addition of necrostatin-1 failed to prevent CX-3543-induced cell death in U2OS cells, suggesting that the observed cell death occurs *via* a mechanism independent of necroptosis (Figure S2D). However, given that U2OS cells lack RIP3, a critical component of the necroptosis pathway, we further examined the potential of CX-3543 to induce necroptosis in Jurkat cells, which are capable of undergoing this form of cell death [33]. To determine the IC50 of CX-3543 in Jurkat cells, we first conducted a cell viability assay, as shown in Figure S2C. In Jurkat cells, necrostatin-1 failed to inhibit cell death triggered by CX-3543, although it effectively reduced cell death induced by the combination treatment of TNFα, QVD-O-Ph, and a SMAC mimetic, AT406 (**Figure 2F**). Together, these findings indicate that CX-3543 induced cell death in both U2OS and Jurkat cells through a mechanism independent of necroptosis.

This indicates that neither apoptosis, ferroptosis, necroptosis nor cathepsins participate in the cell death induced by CX-3543.

### CX-3543 triggers TFEB activation to cope with lysosomal stress

In response to a variety of cellular stresses, TFEB and its homolog TFE3 translocate from the cytosol to the nucleus and trigger transcriptional programs to activate adaptive responses [15,34–36]. TFEB activity is tightly controlled by the mechanistic target of rapamycin complex 1 (mTORC1), which retains TFEB in the cytoplasm by inducing its phosphorylation at 142 and 211 serine residues (**Figure 3A**). While under basal conditions, endogenous TFEB and TFE3 were predominantly localized in the cytoplasm of U2OS cells, exposure to CX-3543 resulted in their nuclear translocation that exceeded even the translocation induced by torin-1, a well-known TFEB-and TFE3-activator (**Figure 3B**). A similar effect was found in HeLa and SaOS-2 cells (Figure S3A). Moreover, we observed that mTOR activity was markedly diminished in a time-dependent manner in response to CX-3543, as indicated by reduced phosphorylation of mTOR itself at residue S2448 and of its substrate p70-S6K at residue S371 (**Figure 3C**, Figure S3B). In line with this result, TFEB was dephosphorylated at residue S211 as early as 30 min after CX-3543 addition (**Figure 3C**). Altogether, these findings suggest that CX-3543-induced mTORC1 inactivation facilitates TFEB activation.

**Figure 3.**
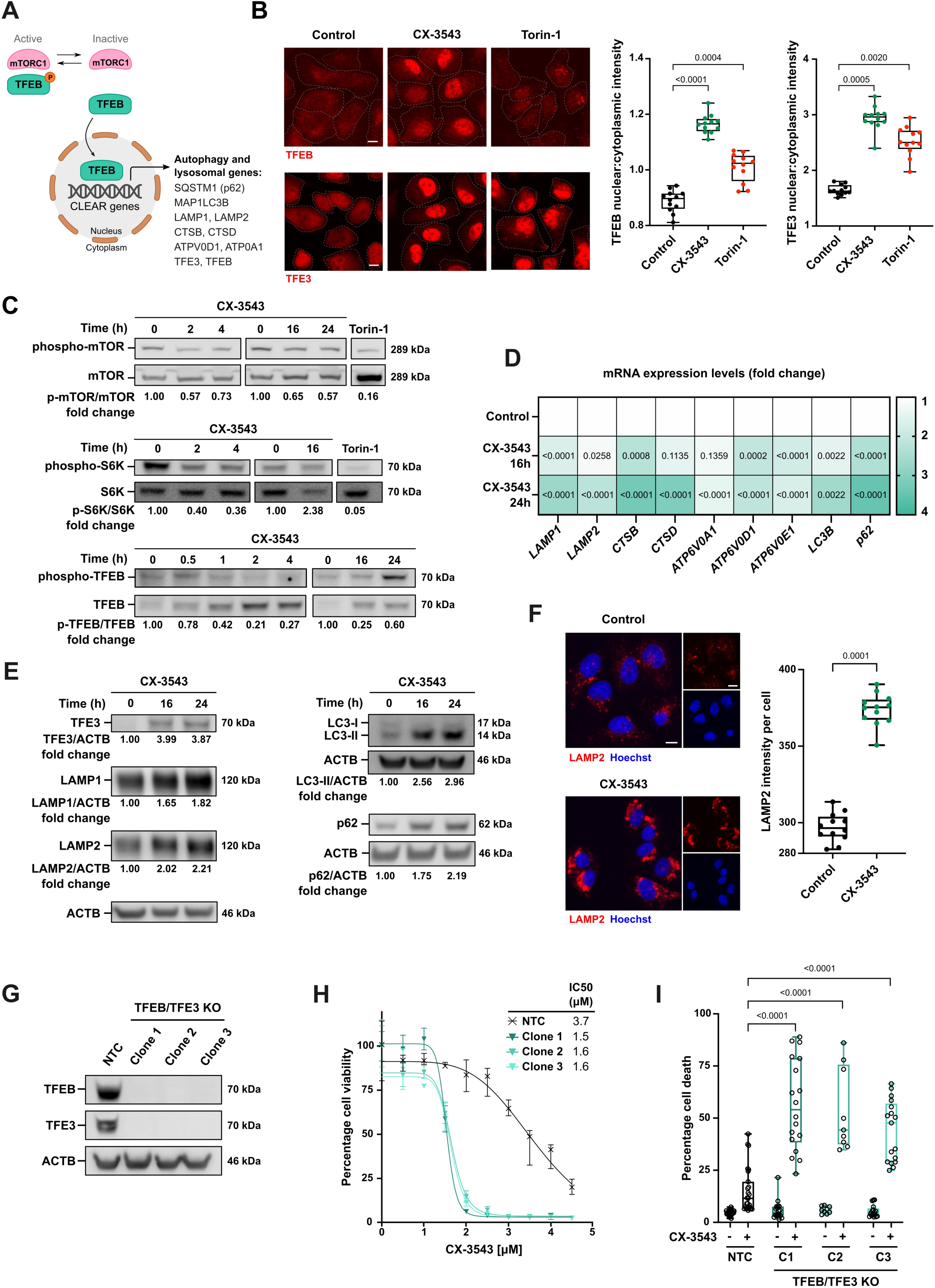
CX-3543 triggers TFEB activation to cope with lysosomal stress. **A)** A simplified diagram illustrates the TFEB pathway. **B)** U2OS cells were treated with either 3 µM CX-3543 or 600 nM torin-1 (a positive control for TFEB and TFE3 nuclear translocation) for 4 h. Post treatment, cells were immunostained with either TFEB or TFE3 followed by HCM analysis. The nuclear-to-cytoplasmic fluorescence intensity ratios of TFEB or TFE3 were measured in at least 2,400 cells per condition. **C)** Western blotting analysis of mTORC1 signaling in U2OS cells treated with either 3 µM CX-3543 or 600 nM torin-1 (a potent inhibitor of mTORC1) at indicated time points. Phosphorylation levels were quantified relative to total protein and normalized to untreated cells. **D)** Real Time-quantitative PCR assessed the expression levels of TFEB target genes in U2OS cells exposed to 3 µM CX-3543 for either 16 h or 24 h. Data were normalized to Peptidyl Propyl Isomerase A (PPIA) housekeeping gene, with the results presented as fold increases relative to untreated cells (N=3). **E)** Immunoblot analysis examined TFEB target gene products in U2OS cells following treatment with 3 µM CX-3543. Band intensity was quantified relative to ACTB/β-actin and normalized to untreated cells. **F)** U2OS cells were treated with 3 µM CX-3543 for 24 h and immunostained for LAMP2. HCM analysis measured the endogenous LAMP2 puncta fluorescence intensity per cell, with a minimum of 2,500 cells analyzed per condition. **G-I)** The role of TFEB and TFE3 in cell death induced by CX-3543 was investigated using Non-Targeting Control (NTC) and *TFEB* and *TFE3* double knock-out (DKO) U2OS cells. **(G)** Western blot analysis of TFEB and TFE3 expression levels. **(H)** Cell viability was assessed by MTT assay after 24 h treatment with CX-3543 (from 0 to 4.5 µM). **(I)** Cell death was evaluated using a Live/Dead Far Red assay after CX-3543 (3 µM) treatment for 24 h (N=4). **Data analysis**: Statistical significance was calculated by means of a robust linear mixed model (**3B and F**), a two-tailed unpaired Mann-Whitney test (**3D**), a non-linear regression test (**3H**) or a 2-coefficients robust linear model (**3I**). Scale bars: 10 μm.

Next, we investigated the impact of CX-3543 on the expression of TFEB target genes, especially those involved in the lysosomal/autophagy pathway [15]. CX-3543 significantly increased the transcription of genes involved in lysosomal and autophagic functions, such as lysosomal associated membrane protein 1 (*LAMP1*) and 2 (*LAMP2*), cathepsin B (*CTSB*), microtubule-associated proteins 1A/1B light chain 3B (*MAP1LC3* best known as *LC3*) and sequestosome-1 (*SQSTM1*, best known as *p62*) (**Figure 3D**). Accordingly, CX-3543 upregulated LAMP1, LAMP2, the active lipidated form of LC3 (LC3-II) and p62 at the protein level (**Figure 3E**). Similarly, immunofluorescence experiments revealed that CX-3543 increased LAMP2 staining intensity (**Figure 3F**), confirming elevated lysosomal biogenesis. However, although a G4-ligand has been reported to control ATG7 expression in neurons, CX-3543 did not significantly affect ATG7 mRNA and protein levels in cancer cells [37] (Figure S3C and D).

We next addressed the functional significance of TFEB activation in cell death induced by CX-3543. Since TFEB and TFE3 exhibit redundant functional cooperativity, we generated U2OS clones stably depleted from both TFEB and TFE3 (*TFEB* and *TFE3* DKO cells) (**Figure 3G**). The cytotoxic effects of CX-3543 were markedly increased in the absence of both TFEB and TFE3, with an IC_50_ dropping from 3.7 to 1.5 µM in *TFEB* and *TFE3* DKO cells (**Figure 3H and I**).

Altogether, these data revealed that CX-3543 induces both TFEB and TFE3 activation associated with the cytoprotective upregulation of several essential genes/proteins involved in the lysosomal/autophagy pathway.

### Regulation of autophagy by CX-3543

The activation of TFEB and TFE3 by CX-3543 raised the question of whether CX-3543 can stimulate autophagy, a lysosomal catabolic process [6,7]. Autophagy occurs in a multistep process [38] that requires the participation of a set of autophagy-related (ATG) proteins, including the autophagy initiation ULK1 complex, the nucleation phosphoinositid-3-kinase class 3 (PI3K3) complex and the elongation and maturation complex (ATG12-ATG5-ATG7). Bafilomycin A1, an inhibitor of V-ATPase, disrupts the final steps of autophagy by preventing lysosomal acidification (**Figure 4A**). Autophagic flux was monitored by following the subcellular relocation of LC3 to autophagosomes in the presence and absence of bafilomycin A1. If autophagy flux is activated, LC3 redistribution to autophagosomes is increased in the presence of bafilomycin A1 [39]. As shown in **Figure 4B**, bafilomycin A1 amplified the number of LC3 puncta in cells exposed to CX-3543 for 16 h. In contrast, no significant effect was observed for those exposed to CX-3543 for 6 h (Figure S4A). The activation of autophagy flux after 16h (but not 6h) of treatment with CX-3543 was further confirmed by immunoblot detection of the lipidated, membrane-associated form of LC3 that was induced by CX-3543, in particular upon addition of bafilomycin A1 (**Figure 4C**, Figure S4B). Altogether, these results indicate that autophagic flux was enhanced after 16 h treatment with CX-3543 coinciding with TFEB- and TFE3-induced transcriptional upregulation of autophagy-relevant genes, both at the mRNA and protein levels 16 and 24 h after CX-3543 addition (**Figure 3D and E**, Figure S4C and D).

**Figure 4.**
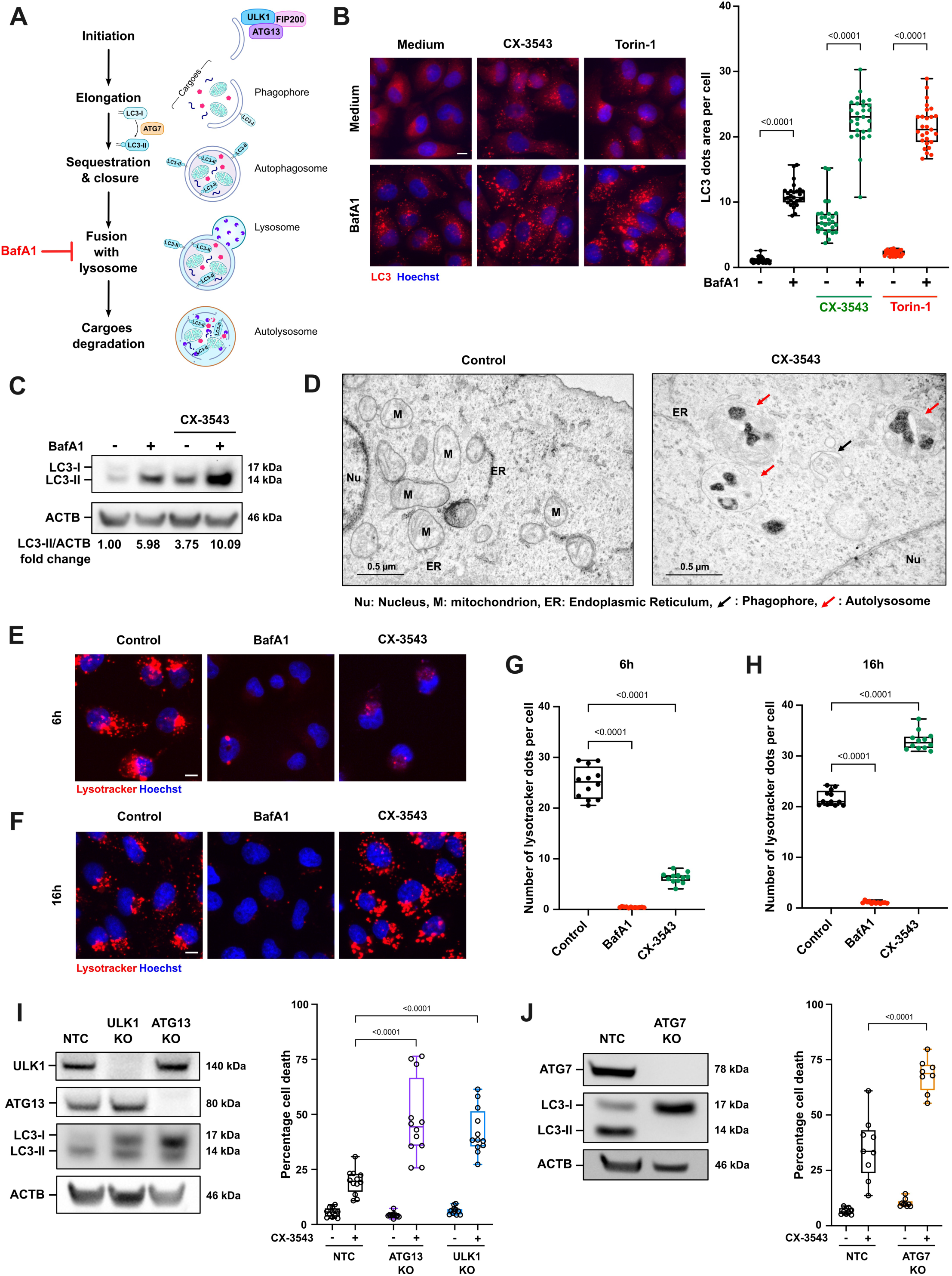
CX-3543 induces the activation of cytoprotective autophagy. **A)** A diagram outlines the key steps of autophagy. **B)** U2OS cells were treated with CX-3543 (3 µM) and 10 nM bafilomycin A1 for 16 h and immunostained for LC3. HCM analysis measured LC3 puncta area per cell, with at least 1,700 cells analyzed per condition. Torin-1 (600 nM, 4 h) served as a positive control for autophagy induction. **C)** Western blot assessed the levels of the lipidated form of LC3 (LC3-II) in U2OS cells treated with either CX-3543 (3 µM, 16 h) or Bafilomycin A1 (10 nM, 16 h). Band intensities were quantified relative to ACTB/β-actin and normalized to untreated cells. **D)** Transmission electron microscopy visualized cellular ultrastructures showing phagophore (black arrows) and autolysosome (red arrows) in U2OS cells treated with CX-3543 (3 µM) for 16h. **E-H)** U2OS cells were treated with 3 µM CX-3543 for either 6 h (**E and G**) or 16 h (**F and H**) before incubation with Lysotracker™ Red DND. Bafilomycin A1 (25 nM, 2 h) served as an inducer of lysosomal disruption. HCM analysis evaluated the number of functional lysosomes per cell, analysing at least 2,400 cells per condition. **I)** The roles of ULK1 and ATG13 in CX-3543-induced cell death were examined using Non-Targeting Control (NTC), *ULK1* KO, and *ATG13* KO U2OS cells. Immunoblotting was used to assess ULK1, ATG13, and LC3 expression levels. Cell death was measured as described in Figure 2B after 24 h treatment with CX-3543 (N=4). **J)** The role of ATG7 in CX-3543-induced cell death was studied in NTC and *ATG7* KO HeLa cells. Immunoblotting evaluated ATG7 and LC3-I/II expression. Cell death was quantified using a Live/Dead Far Red assay after 24 h treatment with CX-3543 (N=3). **Data analysis** Statistical significance was calculated by means of robust linear models: a two-coefficients robust linear mixed model (**4B, I and J**), or a one-coefficient robust linear mixed model (**4G-H**). Scale bars: 10 μm.

Accordingly, transmission electron microscopy revealed that U2OS cells treated with CX-3543 for 16 h displayed a notable increase in the abundance of phagophores and vesicles containing cellular debris (**Figure 4D**). To determine whether the activation of autophagy flux was linked to increased lysosomal biogenesis and function, we utilized the Lysotracker Red dye. U2OS cells treated with CX-3543 exhibited a noticeable decrease in Lysotracker staining after 6 h of treatment compared to untreated cells, followed by a significant increase after 16 h of treatment (**Figure 4E-H**). This biphasic response to CX-3543 might be linked to initial LMP (at 6 h) followed by a subsequent adaptive response (at 16 h) coinciding with the enhancement of autophagic flux.

To address the functional significance of autophagy activation in regulating cell death induced by CX-3543, we used U2OS cells deficient in ULK1 or ATG13 to block the initial step of autophagy (**Figure 4I**), as well as HeLa cells lacking ATG7, in which LC3 lipidation is impaired (**Figure 4J**). As shown in **Figure 4I** and **J**, CX-3543-induced cell death was significantly increased in all autophagy-deficient cells compared to their control counterparts. Together, these results suggest that CX-3543 activates autophagy as an adaptive mechanism to protect cells against cell death.

### Targeting lysosomes enhances the anticancer efficacy of the POL I transcription inhibitors, CX-3543 and CX-5461

We next investigated whether disrupting lysosomal function and integrity would increase cancer cells sensitivity to CX-3543. To test this hypothesis, we used three well-established inhibitors of lysosomal functions: chloroquine and two of its derivatives, Lys05 and DC661.

To evaluate the interaction between CX-3543 and DC661, we measured cell viability and conducted an isobologram analysis utilizing the Harbron’s method [40,41], wherein combination indexes (CI) were computed. CI values calculated for the combinations of CX-3543 and DC661 were consistently <1, indicating a synergistic interaction between both agents (**Figure 5A and B**). Based on these data, we selected concentrations of CX-3543 (2 µM) and DC661 (1 µM) resulting in approximately 40% cell viability loss (Table S2). Moreover, the combination of CX-3543 and chloroquine or its derivatives induced a marked increase in cell death of U2OS cells compared to the individual treatments (**Figure 5C**, Figure S5A and B). A similar synergic effect was observed when HeLa cells were treated with CX-3543 combined with either chloroquine or Lys05 (Figure S5D-F). CX-5461 also significantly increased U2OS cell death when combined with Lys05 or DC661 (**Figure 5D**, Figure S5C). This suggests that lysosomal destabilization enhances the lethal impact of the two CX compounds. To further characterize the cell death induced by the combined treatments, we used the pan-caspase inhibitor Q-VD-OPh. As shown in **Figure 5E and F**, Q-VD-OPh only had a modest effect on cell death induced by the combined treatment. Moreover, CX-3543 plus DC661 did not induce detectable Caspase-3 cleavage as shown by western blotting, suggesting that cell death primarily occurs through an apoptosis-independent mechanism (Figure S5G).

**Figure 5.**
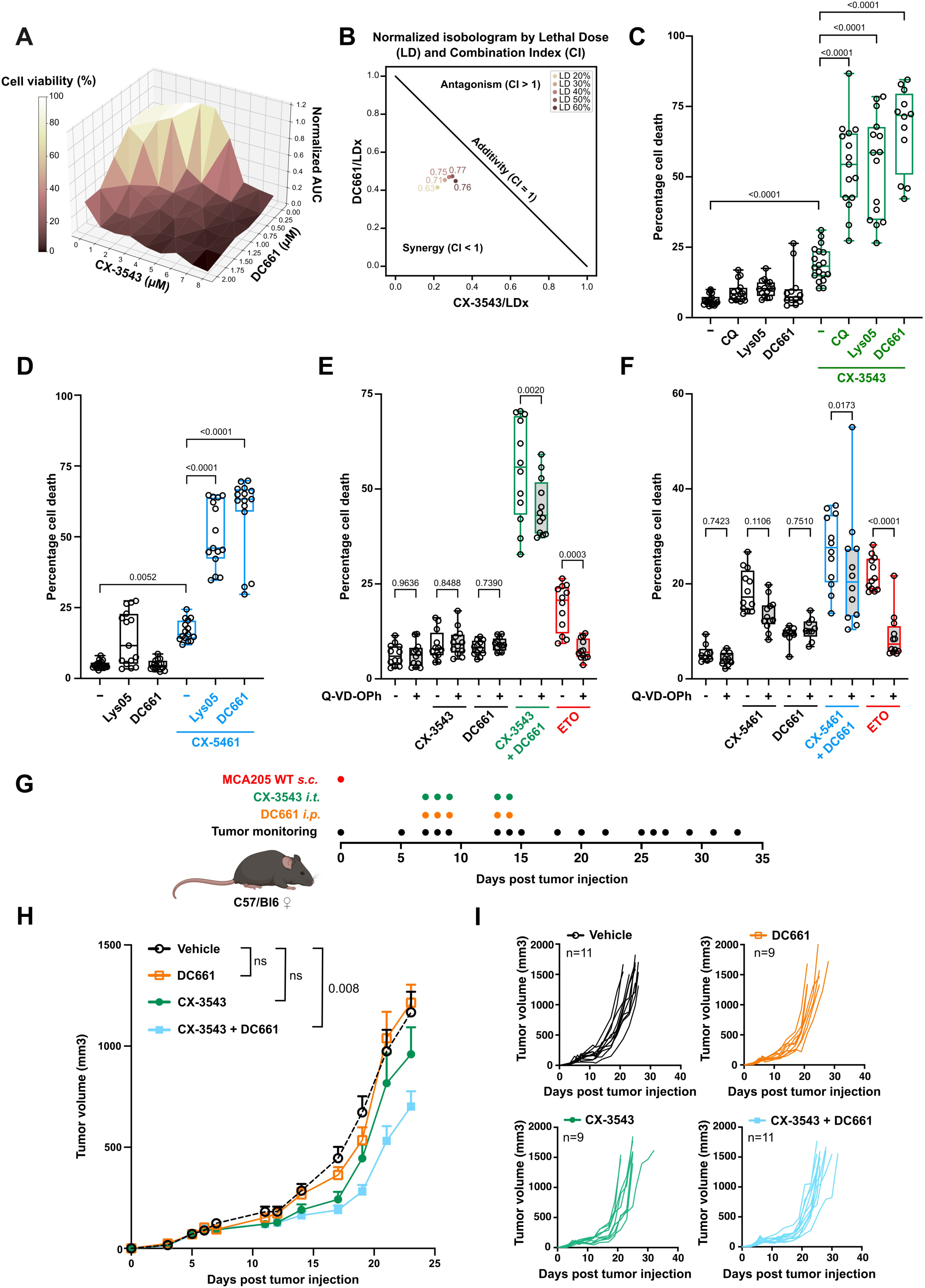
Chloroquine derivatives amplify the anticancer efficacy of POL I transcription inhibitors. **A-B**) U2OS cells treated by CX-3543 (1-8 µM), DC661 (0.4-2 µM), or their combination. Cell viability was then monitored hourly for 24 h using live label-free microscopy. **A)** Area under each dose curve (AUC) was normalized to the time zero and plotted in 3D showing mean response. Cell viability is color-coded from yellow (100%) to brown (0%). **B)** The combination indexes (CI) for CX-3543 and DC661 were calculated for a given lethal dose (LD) using the Harbron method [58]. CI <1, =1, >1 indicate the concentrations at which the combination of drugs was synergistic, additive, and antagonistic, respectively. **C-D)** U2OS cells were treated for 24h with: **(C)** CX-3543 (2 µM) in the presence or absence of chloroquine (15 µM), Lys05 (2 µM) or DC661 (800 nM). **(D)** CX-5461 (3 µM) and the same treatments as above. Cell death was then measured using a Live/Dead Far Red assay (N=4). **E-F)** U2OS cells were initially treated with Q-VD-OPh (40 µM) for 2 h, followed by treatment with either CX-3543 (2 µM) or CX-5461 (3 µM), administrated alone or in combination with DC661 (800 nM). Etoposide (100 µM, 24 h) was utilized as an inducer of apoptotic cell death (N=4). After 24 h post-treatment, cell death was assessed using a Live/Dead Far Red assay. **G-I)** Experimental schedule of implantation and treatment of murine fibrosarcoma in mice. **G)** Murine fibrosarcoma MCA205 cells were inoculated sub-cutaneously into the right flank of immunocompetent C57BL/6 WT female mice. When tumors were palpable, mice received treatments with vehicle PBS/DMSO (n=11 mice per group), 5 mg/kg CX-3543 (intratumoral injections, n=9 mice per group), 3 mg/kg DC661 (intraperitoneal injections, n=9 mice per group) or a combination of the two drugs (n=11 mice per group). Treatment occurred on days 7, 8, 9, 13, and 14 post-tumor inoculation. Tumors were measured three times weekly. **H)** Results represent the mean tumor volume for each group. **I)** Results represent the tumor volume of individual mouse. **Data analysis**: Statistical significance was calculated by means of a 2-coefficients robust linear mixed model (**5C-F**), or a TumGrowth test (https://github.com/kroemerlab) (**5H-I**).

Next, we investigated the effect of CX-3543 plus DC661 in an *in vivo* xenograft immunocompetent tumor model. For this, we established subcutaneous MCA205 fibrosarcoma in immunocompetent mice and monitored tumor growth after treatment with CX-3543 and DC661, alone or in combination (**Figure 5G**). We used minimal doses of each drug to discern a potential synergistic effect. While administration of CX-3543 or DC661 alone had no significant effect, the combined treatment with CX-3543 and DC661 induced a decrease in tumor growth and an increase in survival rates compared to control mice (**Figure 5H and I**, Figure S5H).

Taken together, the data imply that lysosomal destabilization enhances the tumor growth inhibition induced by CX-3543.

### DC661 enhances the lysosomal and nuclear toxicity of CX-3543 and CX-5461

Having established a synergistic effect between chloroquine derivatives and either CX-3543 or CX-5461, we next aimed to understand the underlying mechanism behind it.

We then determined whether chloroquine derivatives sensitize cells to death triggered by CX-3543 or CX-5461 through a mechanism involving LMP induction. This would allow CX-3543 and CX-5461 to enter the nucleolus and subsequently inhibit POL I transcription to cause cell death. To test this hypothesis, we first measured LMP by means of the LGALS3-mCherry biosensor in cells treated by either CX-3543 or CX-5461 in the presence and absence of DC661. We used the same sublethal doses for each compound as in the cell death experiments (**Figure 5C and D**). The combined treatment with DC661 and CX-3543 significantly increased the number of LGALS3 puncta per cell compared to CX-3543 alone. Similarly, DC661 markedly enhanced LGALS3 puncta formation induced by CX-5461 (**Figure 6A**). In the same vein, chloroquine and Lys05 significantly increased LMP induction by CX-3543 (Figure S6A). Finally, combinations of DC661 with either CX-3543 or CX-5461 resulted in a significant reduction in lysosome function compared to each drug alone (**Figure 6B**). Altogether, these results indicate that DC661 exacerbates lysosomal membrane damage triggered by either CX-3543 or CX-5461.

**Figure 6.**
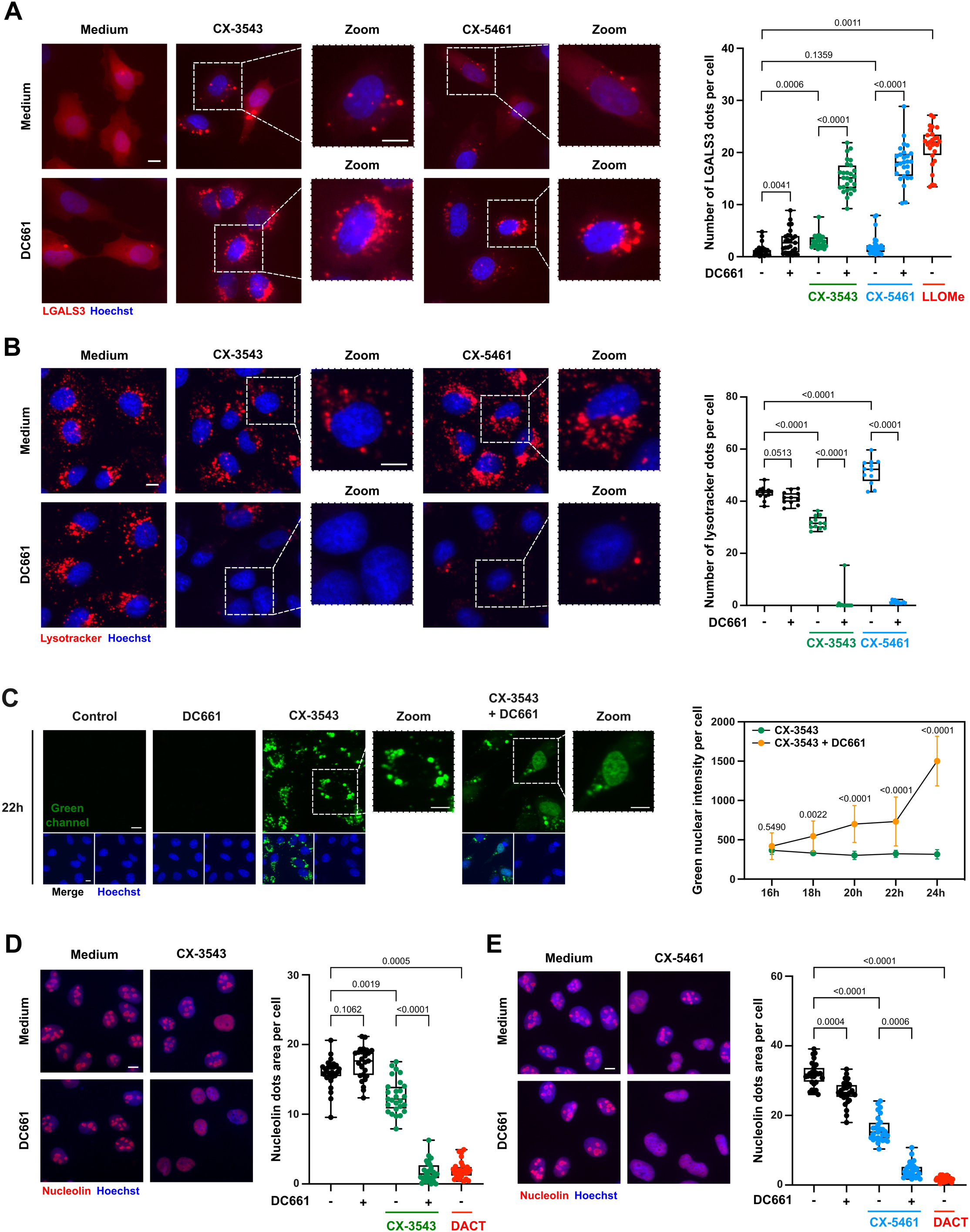
Chloroquine derivatives potentiate lysosomal damage and nucleolar disruption induced by CX-3543 or CX-5461. **A)** U2OS cells stably expressing LGALS3-mCherry were treated with either CX-3543 (2 µM) or CX-5461 (3 µM) alone or in combination with DC661 (800 nM) for 6 h. HCM analysis was performed to evaluate the number of LGALS3 puncta per cell with a minimum of 1,700 cells analyzed per condition. LLOMe (1 mM, 1 h) was used as a positive control for LMP induction. **B)** U2OS cells were treated with CX-3543 (2 µM) or CX-5461 (3 µM) alone or in combination with DC661 (800 nM) for 16 h before Lysotracker™ Red DND incubation. HCM was performed to evaluate the number of functional lysosomes per cell, with a minimum of 2,400 cells analyzed per condition. **C)** To visualize the nuclear translocation of CX-3543, U2OS cells were treated with CX-3543 (2 µM) alone or in combination with DC661 (800 nM) from 24 to 16 h. HCM analysis was performed to evaluate the green nuclear fluorescence intensity of CX-3543 per cell, with a minimum of 1,200 cells analyzed per condition. **D-E)** U2OS cells were treated with CX-3543 (2 µM) **(D)** or CX-5461 (1 µM) **(E)** alone or in combination with DC661 (800 nM) for 16 h and then immunostained with Nucleolin. HCM analysis was performed to evaluate the area of Nucleolin puncta per cell, with a minimum of 1,700 cells analyzed per condition. Dactinomycin (1 µM, 2 h) was used as a positive control for nucleolar disruption. **Data analysis**: Statistical significance was calculated by means of a two-coefficients robust linear mixed model (**6B and C**) followed by a one-coefficient robust linear mixed model (**6A, D, E**). Scale bars: 10 μm.

Next, we determined whether DC661 would facilitate the preferential accumulation of CX-3543 in nuclei rather than in lysosomes. For this, we monitored the blue light-elicited green fluorescence of CX-3543 in the absence or presence of DC661. As shown in **Figure 6C**, while CX-3543 alone was withhold in the cytoplasm during the time course of treatment (16 h - 24 h), the addition of DC661 allowed its relocation into the nuclear compartment after 18 h. This time frame is consistent with the massive LMP induction in cells treated with CX-3543 plus DC661. To further confirm our hypothesis, we next evaluated nucleolar disruption in cells treated with CX-3543 or CX-5461 alone or in combination with DC661. As shown in **Figure 6D and E**, the combined treatments significantly reduced nucleolar Nucleolin staining compared to exposure to CX-3543 or CX-5461 alone. Moreover, DC661 plus CX-3543 or DC661 plus CX-5461 notably reduced RNA transcription as evidenced by the 5-ethynyl uridine (5-EU) incorporation assay [29] (Figure S6C).

Altogether, these results support the idea that DC661-induced LMP boosts nucleolar disruption and RNA transcription inhibition triggered by CX-3543 or CX-5461.

### Blue light excitation enhances CX-3543-induced cytotoxicity

Given that CX-3543 emits a green fluorescence upon blue light excitation, we wondered whether this agent would photosensitize cells to blue light or vice versa. To test this hypothesis, we used LGALS3-mCherry expressing cells treated with CX-3543 and performed video microscopy with and without blue light excitation. As shown in **Figure 7A** and **C**, illumination with blue light resulted in the early induction of LMP, as indicated by the formation of LGALS3 puncta. The early LMP induction under blue light exposure coincided with a rapid relocation of CX-3543 into the nucleus (**Figure 7A and D**). Furthermore, blue light excitation accelerated the loss of cell viability caused by CX-3543 treatment (**Figure 7B and E**).

**Figure 7.**
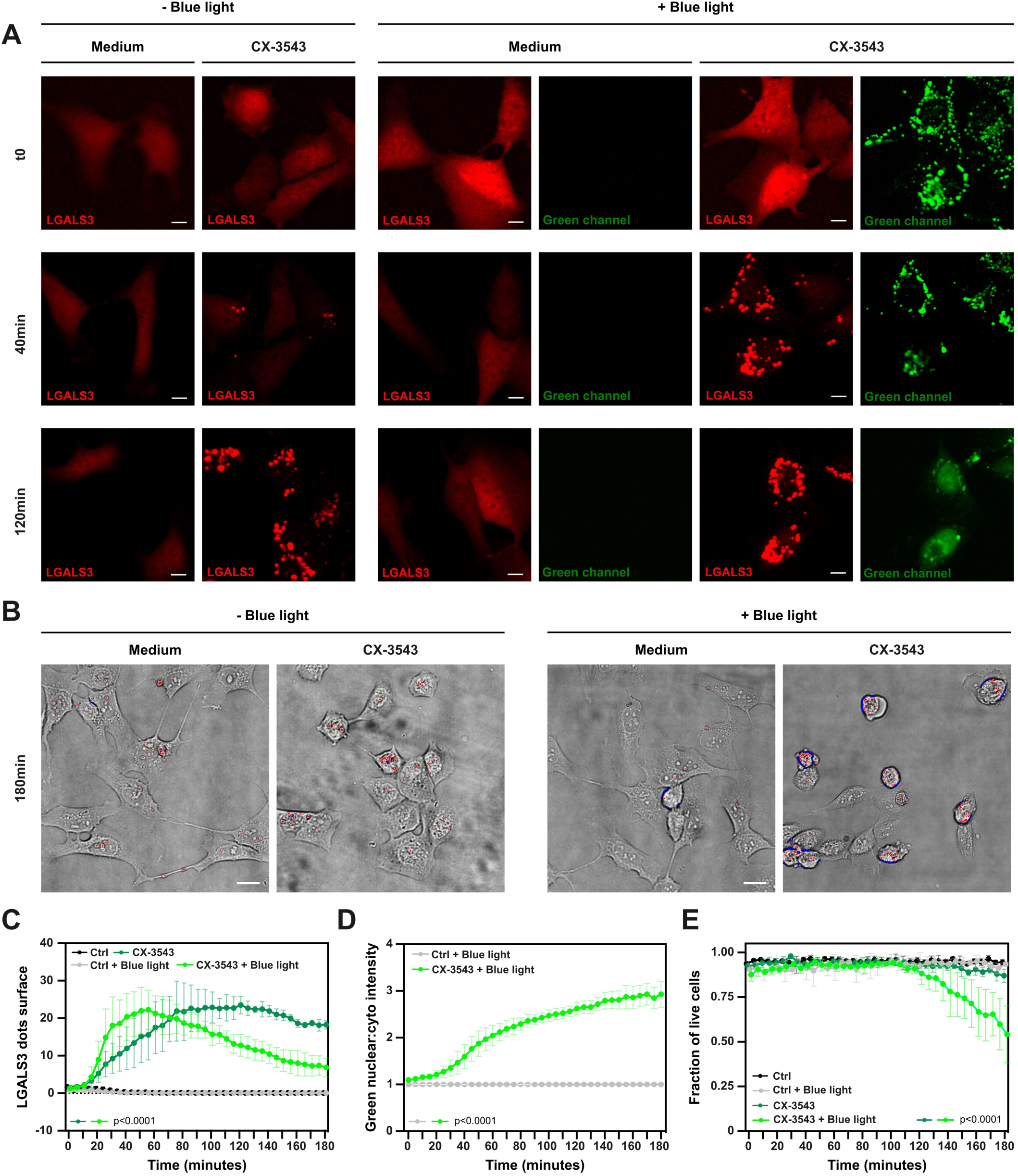
Blue light excitation enhances CX-3543 cytotoxicity by facilitating its release from lysosomes. **A-E)** U2OS LGALS3-mCherry cells were treated with CX-3543 (3 µM) and exposed or not to blue laser illumination (470 nm). Dactinomycin (1 µM) served as a potent inhibitor of POL I transcription. Images were acquired every 5 min for 3 h using a 20X objective. **A)** Representative sequential fluorescence images of LGALS3 puncta formation (red signal), CX-3543 subcellular localization (green signal), and cell viability (Transmitted light 100) are shown. The quantifications represent **C)** the LGALS3 puncta surface, **D)** the green nuclear:cytoplasmic intensity ratio **E)** cell viability through a model trained with transmitted light images as described in the materials and methods section. The color code indicates the degree of cell refringence using the look up table (HiLo) tool in ImageJ. Red color depicts the most refringent cells, while blue color indicates the least refringent cells. **Data analysis**: Statistical significance was calculated by means of the following models: **7C)** a quadratic regression model for the 4 curves, **7D)** a linear regression model for the 2 curves, **7E)** two linear regression models, one before and one after 110 min. See the materials and methods section for more detailed information. Scale bars: 10 μm.

Collectively, these results suggest that blue light excitation sensitizes cancer cells to CX-3543 by inducing LMP **(Figure 8)**.

**Figure 8.**
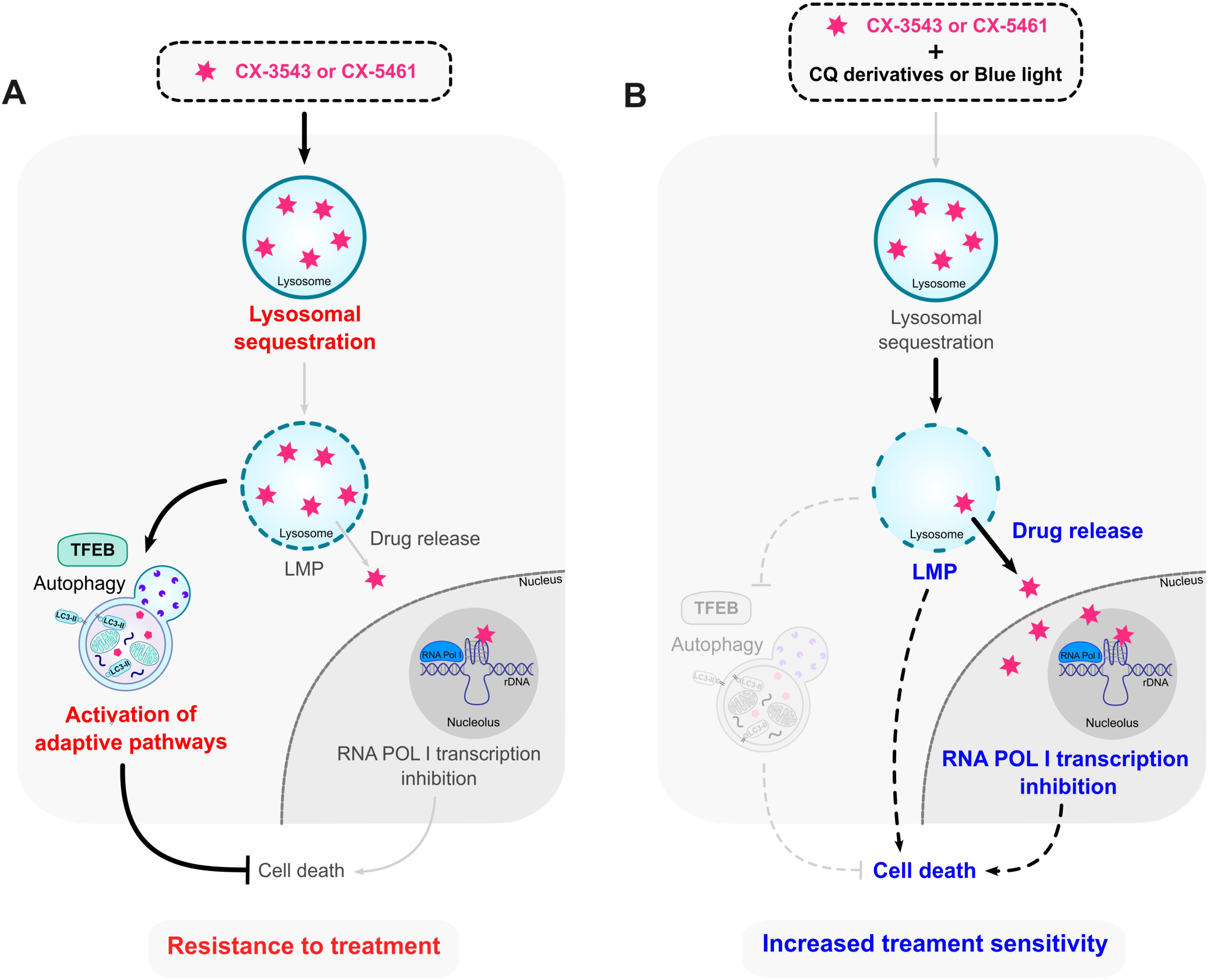
Targeting lysosomal membranes amplifies POL I transcription inhibition and cell death induced by CX-3543 and CX-5461. **A)** The POL I transcription inhibitor, CX-3543 is sequestrated in lysosomes, where it induces lysosomal membrane permeabilization (LMP). As a result, TFEB and autophagy, two adaptive stress responses, are activated to cope with lysosomal stress. This results in increased lysosomal biogenesis, leading to elevated lysosomal sequestration of the compound, thereby impeding its inhibitory effect on POL I transcription. The observed lysosomal effects may play a role in conferring resistance to cancer cells against these treatments. **B)** The combined treatment of POL I transcription inhibitors, CX-3543 and CX-5461, and either chloroquine derivatives or blue light excitation (in the case of CX-3543) yields a marked increase in LMP induction while concurrently suppressing the autophagy pathway. This elevated LMP facilitates the release of the compounds from the lysosomes, allowing the drugs to reach their nuclear targets to inhibit effectively POL I transcription. These combined treatments could represent a promising strategy to overcome resistance to POL I transcription inhibitors such as CX-3543 and CX-5461 and sensitize cancer cells to death.

## Discussion

Elevated ribosomal RNA synthesis by RNA Polymerase I (POL I) in the nucleolus is associated with abnormal cancer cell proliferation and poor prognostic outcomes across various cancers [42,43]. Targeting RNA Polymerase I thus emerges as an attractive antineoplastic strategy [43]. However, only a few RNA POL I transcription inhibitors have shown promising results in preclinical models and clinical trials. This limitation is due to concerns about toxicity, potential side effects, poor bioavailability, and resistance mechanisms [44,24,23,45]. Here, we reveal unanticipated lysosomal-related mechanisms hampering the anticancer effects of two specific POL I transcription inhibitors. We provide evidence that dual targeting of lysosomal membranes and POL I inhibition led to significant cell death through a mechanism that is largely independent of apoptosis. This combination not only overcomes resistance to POL I inhibitors but also mitigates their toxicity by reducing the doses that must be administrated. This combination treatment is particularly pertinent for tumors that resist therapy-induced apoptosis.

The sequestration of drugs within lysosomes contributes to resistance to treatment as it prevents the drugs from reaching their intended targets [11,46,47]. Numerous DNA-binding molecules used in clinical settings (e.g., daunorubicin, doxorubicin, and mitoxantrone) undergo lysosomal sequestration, reducing their effectiveness and contributing to therapeutic resistance [46]. Generally, these drugs display the characteristics of lysosomotropic agents, with physicochemical properties allowing them to become sequestered in lysosomes [48]. To counteract this drug resistance, agents that increase lysosomal pH or trigger LMP have been used [18]. Our *in-silico* studies suggested that CX-3543 and CX-5461 (but not dactinomycin, another POL I inhibitor) share lysosomotropism. This supports the idea that the observed effect on LMP is due to these properties rather than POL I inhibition. Moreover, amplifying LMP induction by DC661 or blue light exposure enables CX-3543 to enter the nucleus and subsequently inhibit POL I transcription, thereby amplifying its cytotoxic potential. This supports the benefit of targeting the lysosomal membrane to reverse CX-3543 lysosomal sequestration and increase the sensitivity of cancer cells to this compound.

Autophagy is a catabolic lysosomal pathway activated in response to various stressful conditions (e.g., nutrient starvation, hypoxia, organellar and DNA damage) to ensure cellular adaptation and survival [49]. Protective autophagy is observed in response to various anti-cancer agents and contributes to resistance to therapy [8,50]. Here, we showed that CX-3543 elicits a homeostatic regulation of autophagy. After short-time (6h) exposure to CX-3543, autophagy is impaired, presumably due to a partial loss of lysosomal function. However, upon long-term (16 – 24 h) treatment, CX-3543 reactivates lysosomal and autophagy activities through TFEB- and TFE3-mediated mechanisms. Accordingly, two lysosomal membrane proteins, LAMP1 and LAMP2, were both upregulated after 16 hours of CX-3543 treatment, supporting the induction of lysosomal biogenesis, which may help maintain lysosomal function. Hence, the suppression of TFEB and TFE3 or autophagy-relevant genes elevates the susceptibility of cancer cells to CX-3543-induced cell death, demonstrating autophagy contribution to resistance mechanisms. Chloroquine derivatives impede autophagy and lysophagy by blocking the fusion of autophagosomes with lysosome [51]. Hence, they may sensitize cells to CX-3543 not only by boosting LMP but also by disrupting autophagy/lysophagy.

Besides TFEB activation, other potential mechanisms may explain lysosome/autophagy pathway activation when POL I transcription is inhibited. In fact, the nucleolar protein Nucleophosmin (NPM) contributes to autophagy activation in response to POL I inhibition but is dispensable for autophagy induction by nutrient starvation [52]. Moreover, the induction of nucleolar stress by various means triggers the upregulation of autophagy-related 7 (ATG7) [53]. Thus, nucleolar stress resulting from POL I inhibition may occur upstream of the activation of autophagy. However, autophagy may also affect ribosomal RNA synthesis. Indeed, autophagy-deficient cells exhibit much higher ribosomal DNA transcription due to the autophagy receptor SQSTM1/p62 accumulation [54]. These findings suggest that the regulation of autophagy and nucleolar stress are caused by POL I inhibition. Future studies should disentangle the crosstalk between both phenomena.

Altogether, we uncovered an unexpected lysosome-related mechanism that attenuates the cytotoxic effects of two POL I inhibitors, CX-3543 and CX-5461. This mechanism entails lysosomal drug sequestration and the activation of the adaptive TFEB/autophagy pathway. Our finding supports the synergistic use of these POL I inhibitors with lysosomal targeting agents or, in the case of CX-3543, with photodynamic therapy (Figure 8). These combination treatments may overcome resistance to these POL I inhibitors and minimize toxicity. Additionally, photodynamic therapy can precisely circumscribe drug effects to the illuminated tissue, enhancing treatment specificity.

## Materials and methods

### Cell culture

The HeLa (human cervical carcinoma), U2OS (human osteosarcoma), Saos-2 (human osteosarcoma) and Jurkat (human leukemic T cell lymphoblast) cell lines were purchased from the American Type Culture Collection (respectively, CCL-2 ™, HTB-96 ™, HTB-85 ™, TIB-152 ™). MCA205 (mouse fibrosarcoma) cells were obtained from Merck (Sigma-Aldrich, SCC173). U2OS cells stably expressing LGALS3-mCherry, *ULK1* KO U2OS cells and *ATG13* KO U2OS cells are generous gifts from Dr. Harald Wodrich (CNRS UMR5234, Bordeaux, France). *ATG7* KO HeLa cells were previously generated in our laboratory [19]. Non-Targeting Control (NTC), *TFEB* and *TFE3* DKO U2OS cells were generated as described in the Lentiviral vector production and cell transduction section.

HeLa, U2OS, and Saos-2 cell lines were all cultured in Dulbecco’s modified Eagle’s medium (DMEM; Thermo Fisher Scientific, 41966029) culture medium supplemented with 10% Fetal Bovine Serum (Sigma-Aldrich, F7524) and HEPES Buffer Solution (10 mM) (Thermo Fisher Scientific, 15630-056). MCA205 cells were grown in Roswell Park Memorial Institute (RPMI) (RPMI; Thermo Fisher Scientific, 61870044) culture medium supplemented with 10% fetal bovine serum and 10 mM HEPES buffer. For U2OS cells stably expressing LGALS3-mCherry, 200 μg/mL Hygromycin B (Invitrogen, 10687010) was added in cell culture medium. Hygromycin B was removed 24 h before each experiment. All the cell lines were cultivated at 37^◦^C in a humidified incubator with 5% CO_2_ and were free from mycoplasma contamination (PCR detection test).

### Lentiviral vector production and cell transduction

The sequence guide RNA were designed using CRISPOR algorithm [55] and detailed in Table S3. Alt-R®-crRNA corresponding to target sequences were purchased from Integrated DNA Technologies (IDT) as well as human crRNA negative control and resuspended to 100 µM in Tris-EDTA buffer (IDT) (Integrated DNA Technologies, 11-01-02-02). They were then equally mixed with 100 µM Alt-R®-tracrRNA (IDT), annealed by heating for 5 min at 95°C, and cooled to room temperature (RT). This two-part guide RNA was mixed in a 1.2-fold excess with 4.5 pmol of Alt-R® S.p-Cas9HIFIv3 (IDT) (Integrated DNA Technologies, 1081061) and transfected into cells using Lipofectamine CRISPRmax reagent (Thermo Fisher Scientific, CMAX00001) following manufacturer instructions. At 2–3 days post-transfection, cells were trypsinized, and half of them were cultured while the other half was pelleted, lysed, and used as PCR template using Phire Tissue Direct PCR Master Mix (Thermo Fisher Scientific, F170L) by using specific primers listed in Table S3. The sequencing of PCR products was done by Eurofins Genomics, and Sanger data were used to quantify Indels reflecting gene knock-out (KO) using the DECODR algorithm [56] or ICE algorithm [57]. Cells were seeded at 0.5 cell/well in 96 well plates and allowed to expand to obtain cellular clones, which were then characterized by Sanger sequencing and western blot analysis. U2OS *TFEB* and *TFE3* DKO clones were obtained using TFEB1-G1 gRNA and TFE3-G2 gRNA sequences by two consecutive transfections.

### Antibodies and reagents

#### Reagents

- Bafilomycin A1 (Sigma-Aldrich, B1793).
- Chloroquine diphosphate salt (Sigma-Aldrich, C6628).
- CX-3543 (MedChemExpress, HY-14776).
- CX-5461 (MedChemExpress, HY-13323).
- Dactinomycin (Sigma-Aldrich, A1410).
- DC661 (MedChemExpress, HY-111621).
- E64d (MedChemExpress, HY-100229).
- Etoposide (MedChemExpress, HY-13629).
- Erastin (Cayman Chemical, 17754).
- Ferrostatin-1 (Calbiochem, 341494).
- Necrostatin-1 (Sigma-Aldrich, N9037).
- Xevinapant (AT406) (Selleckchem, S2754).
- Human TNF-alpha (ThermoFisher Scientific, 300-01A-50UG).
- Hoechst 33342 (Thermo Fisher Scientific, H3570).
- L-Leucyl-L-Leucine methyl ester hydrobromide (LLOMe) (Leu-Leu-methyl ester hydrobromide; Sigma-Aldrich, L7393).
- Lys05 (Sigma-Aldrich, SML2097).
- Pepstatin A methyl ester (EMD Millipore, 516485).
- QV-D-Oph (MedChemExpress, HY-12305).
- Torin-1 (Tocris Bioscience, 4247).

#### Antibodies

- Actin HRP-conjugated (Abcam, ab49900).
- ATG7 rabbit (Cell Signaling Technology, 2631).
- ATG13 rabbit (Sigma-Aldrich, SAB4200100).
- Alexa Fluor 568-conjugated Goat anti-rabbit (Thermo Fisher Scientific, A11036).
- Alexa Fluor 568-conjugated Goat anti-mouse (Thermo Fisher Scientific, A1103).
- Cathepsin B rabbit (Cell Signaling Technology, 31718).
- Cleaved Caspase-3 rabbit (Cell Signaling Technology, 9661).
- LAMP1 rabbit (Cell Signaling Technology, 9091).
- LAMP2 mouse (Santa Cruz Biotechnology, 18822).
- LC3 rabbit (MBL International Corporation, PM036) for immunofluorescence.
- LC3 rabbit (Cell Signaling Technology, 27755) for western blot.
- LGALS3/Galectin 3 rabbit (Cell Signaling Technology, 87985).
- Goat anti-rabbit (Southern Biotech, 4050-05).
- Goat anti-mouse (Southern Biotech, 1031-05).
- Nucleolin rabbit (Cell Signaling Technology, 14574).
- phospho-mTOR Ser2448 rabbit (Cell Signaling Technology, 5536).
- mTOR rabbit (Cell Signaling Technology, 2983).
- phospho-TFEB S211 rabbit (Cell Signaling Technology, 37681).
- TFEB rabbit (Cell Signaling Technology, 4240).
- TFE3 rabbit (Cell Signaling Technology, 14779).
- S6K rabbit (Cell Signaling Technology, 9202).
- phospho-S6K S371 rabbit (Cell Signaling Technology, 9208).
- phospho-S6K T389 rabbit (Cell Signaling Technology, 9205).
- SQSMT1/p62 mouse (BD Biosciences, 610832).
- ULK1 rabbit (Cell Signaling Technology, 8054).

### Cell viability assay

The MTT assay was used to monitor cell viability after drug treatment. Cells were seeded in a 96-well plate (5 × 10^3^ cells/well). The following day, cells were treated with indicated drugs for 24h. After 24 h of treatment, Thiazolyl Blue Tetrazolium (0.5 mg/ml) in PBS (Thermo Fisher Scientific, 10010-015) was added to each well for 3 h incubation at 37^◦^C. The supernatant of cells was then removed, and 50 μL of DMSO was added per well. The absorbance in each well was measured at 570 nm using an absorbance VICTOR® Nivo™ Plate Reader (Perkin Elmer). The IC50 of the drugs was determined using a non-linear regression curve within GraphPad Prism V10.1.1 software. When indicated, isobologram analysis was conducted to determine the combination index (CI) for drug combinations using the Harbron method [58].

### Cell death assessment by flow cytometry

Cell death was determined by measuring the fluorescence of the dye reacting with free amines both at the surface and inside the cells, using the LIVE/DEAD™ Fixable Far Red Dead Cell Stain Kit (Thermo Fisher Scientific, L10120). 5,000 cells were seeded in 96-well plates and treated 24 h later. Where indicated, cells were treated with inhibitors targeting various cell death pathways 2 h prior to drug treatment. After 24h treatment, supernatant and attached cells were collected and pelleted at 500G for 5 min at 4^◦^C. Cells from each well were then resuspended in 100 μl PBS plus 1:200 reconstituted fluorescent reactive dye and incubated on ice in 96-well round bottom plates for 30 min at 4^◦^C, in the dark. Cells were then analyzed by flow cytometry on a MACSQuant cytometer (Miltenyi Biotec) using emission filters appropriate for the LiveDead Far Red probe (laser 635 nm, filter 655-730, channel R1). For each biological replicate, 10,000 cells were first gated on a plot of FSC-A versus SSC-A to eliminate debris. Single cells were then selected using a gating FSC-H versus FSC-W. Then, the percentage of live and dead cells was obtained with the R1 channel (exc: 633/635 nm, em: 665 nm) gating. Data were then analyzed using the Flow-Jo software.

### Immunoblotting

Cells were harvested, pelleted at 500 x g in cold PBS and subsequently lysed. Lysis buffer consisted of 10 mM Tris, pH 7.4, 1% SDS (Sigma-Aldrich, 05030), Complete Mini EDTA free Protease Inhibitor (Roche, 11836170001), and PhosphoSTOP Phosphatase Inhibitor (Roche, 04906837001). The lysed cells were then treated with Benzonase endonuclease (EMD Millipore, 71205) for 5 min at room temperature. The protein concentration in each condition was measured with a BCA protein assay kit (Thermo Fisher Scientific, 23225) using bovine serum albumin (BSA) as standard. Samples (15 μg proteins) were then mixed with a mix of NuPAGE™ LDS Sample buffer (4X) (Thermo Fisher Scientific, NP0008) and NuPAGE™ Sample Reducing Agent (10X) (Thermo Fisher Scientific, NP0009), and heated for 10 min at 65°C before sample loading on NuPAGE™ 4% to 12 % Bis Tris protein gels (Thermo Fisher Scientific). After electrophoresis, the proteins were transferred to a nitrocellulose membrane (Cytiva, 10600001). The blots were then incubated with specific primary antibodies according to the manufacturer’s instructions and followed by an appropriate horseradish peroxidase-conjugated secondary antibody. Immunostained proteins were detected with the enhanced chemiluminescence (ECL) system thanks to the iBright 1500 camera (Invitrogen). ACTB/β-Actin was used as a protein loading control. The densitometry quantification was performed using the ImageJ software.

### Detection of endogenous proteins by immunofluorescence assay

Cells were plated in 96-well plates (8,000 cells/well for up to 8 h treatment and 5,000 cells/well for 16 to 24 h treatment). At the end of the treatment, cells were washed with PBS and fixed with a solution containing PBS 4% paraformaldehyde plus 10 µg/ml Hoechst 33342 for 20 min in the dark, at room temperature. Depending on the protocol, cells underwent permeabilization with PBS containing 3% BSA (Euromedex, 04-100-812-E) and either 0.3% Triton X-100 solution (Sigma-Aldrich, 93443) (for immunostaining of nuclear proteins such as TFEB or Nucleolin) for 10 min or 250 µg/ml Digitonin for 5 min (for immunostaining of membrane-associated proteins such as LC3, LAMP2, and LGALS3). After several washes with PBS, cells were blocked in PBS containing 5% Fetal Bovin Serum to reduce nonspecific antibody binding, for 30 min, before incubation with the primary antibody in PBS and 3% BSA for 2h at room temperature or overnight at 4°C, in the dark. After PBS wash, a secondary antibody prepared in PBS supplemented with 3% BSA was then incubated for 1 h at room temperature, in the dark. After several washes with PBS, the plates were sealed with aluminum microplate sealing tapes (Thermo Fisher Scientific). The images were then acquired using an ImageXpress® Micro Confocal High-Content Imaging System (Molecular Devices) utilizing either the 20X or the 40X objective depending on the readout of the experiments. When the 20X and 40X objectives were used, approximately 60 or 200 cells were scored for each image, respectively.

### Lysosomal membrane permeabilization (LMP) detection using Galectin 3 marker

The lysosomal membrane permeabilization (LMP) was determined by using the specificity of LGALS3 protein, which transitions from a diffuse pattern under normal conditions to a punctuated one upon LMP induction [9]. For these experiments, we utilized U2OS cells expressing LGALS3-mCherry. We also evaluated the formation of endogenous LGALS3 puncta using an immunofluorescence assay as described above. The images were acquired using either the 20X or 40X objective through high-content image microscopy with automated image acquisition (Molecular Devices).

### Fluorescence Image analysis with the Custom Module Editor from MetaXpress

Image segmentation was automated using the Custom Module Editor from MetaXpress Software (Molecular Devices). The primary nuclear region of interest (ROI) was created by masking the nucleus for cell count (and to quantify the fluorescence intensity of nuclear targets). A secondary cytoplasmic ROI was used to measure the fluorescence intensity of the cytoplasmic target and quantify the punctate objects. The masks obtained were further applied to the original fluorescent images to extract relevant measurements, including fluorescence intensity, dot counts/area (for analysis of LC3 and LGALS3 puncta), nucleo-cytoplasmic localization (for TFEB nuclear translocation), and cell count. Data were normalized after the exclusion of debris and dead cells.

### Live-cell microscopy

For monitoring, LMP, drug release from lysosomes, cell death, cells cultured in complete growth medium were subjected to live-cell imaging microscopy. Sequential acquisitions were performed for 3 h, captured in quadruplicates every 5 min, for each treatment condition. Acquisitions were performed on an ImageXpress HT.ai (Molecular Devices) equipped with a CFI PlanApo Lambda 20X objective (Nikon) and appropriate filter sets. Transmitted Light source was used for capturing brightfield images, allowing to compute cell and nuclear masks after applying a pretrained neural-network model (NNM) for semantic segmentation [59]. The TexasRed channel was used to detect LGALS3-mCherry signal. FITC channel was used after blue-laser excitation of CX molecules and collection of green fluorescence. Micrographs were processed using custom R scripts embedded with EBImage [60] MetaxpR and MorphR packages (https://github.com/kroemerlab). Briefly, ‘dots’ were detected after applying a top hat filter with an appropriate kernel to cell images, thereafter subjected to adaptive thresholding. Cells were then classified as “dead” or “alive” based on their circularity, and cellular features (number/surface of punctate, fluorescence intensity) were computed on living cells. The look up table (HiLo) tool in ImageJ was used to show the degree of cell refringence. Last, single cell results were aggregated on an image basis by computing features mean, and resulting quadruplicate values were used to calculate median and median absolute deviation (mad) for each condition.

### Assessment of lysosomal integrity using LysoTracker™ dye

Lysosomal integrity was determined using LysoTracker™ Red DND-99 dye (Invitrogen, L7528). Cells were plated in 96-well plates (8,000 cells/well). At the end of the treatment, LysoTracker™ Red DND-99 probe (150 nM) and Hoechst 33342 10 µg/ml were added to the culture medium for 30 min at 37°C. Subsequently, cells were washed twice with PBS and then incubated with complete medium before imaging using high-content image microscopy with automated image acquisition (Molecular Devices).

### Visualization of lysosomes or mitochondria with LysoTracker™ and MitoTracker™ dyes

Lysosomes and mitochondria visualization was performed using respectively LysoTracker™ Red DND-99 dye (Invitrogen, L7528) and MitoTracker™ Red FM dye (Invitrogen, M22425). Cells were plated in 96-well plates (8,000 cells/well) for 16 h or (5,000 cells/well) for 8 h of treatment. At the end of the treatment, LysoTracker™ or MitoTracker™ probes (150 nM) plus Hoechst 33342 10 µg/ml were added to the culture medium for 30 min at 37°C. Cells were then washed twice with PBS before acquisition using high-content image microscopy with automated image acquisition (Molecular Devices). We then utilized ImageJ software to extract the profile intensity distribution of the blue (Hoechst), red (Lysotracker or Mitotracker), and green (CX-3543) channels from fluorescence images. The obtained values were then used to compute a Pearson Correlation Coefficient within the GraphPad Prism V10.1.1 software (GraphPad Software Inc.).

### Assessment of transcription using Click-iT™ RNA Alexa Fluor™ 594 assay

Transcription of DNA to RNA was measured using the two-step commercial kit Click-iT™ RNA Alexa Fluor™ 594 (Thermo Fisher Scientific), which ligates 5-ethynyl uridine (EU), an alkyne-modified nucleoside capable of being integrated into recently synthesized RNA, to an azide-containing dye, hence allowing the detection of RNA transcription.

Cells were plated into 384-well dark Greiner plates (2,000 cells/condition). One hour before the end of treatment, cells were incubated with 1 mM EU. Cells were then fixed with PBS 4% paraformaldehyde plus 10 µg/ml Hoechst 33342 for 20 min in the dark, at room temperature. Cells were then permeabilized with 0.1% TritonX-100 for 10 min. After the permeabilization solution was discarded, 20 μL per well of the reaction mixture containing the Alexa Fluor®-594-conjugated azide was added to each well. The plate was then incubated for 2 h in the dark at room temperature.

Subsequently, the plate was washed with PBS and sealed with aluminum sealing tapes before acquisition on the Molecular Device plate reader (Molecular Devices).

### Transmission electron microscopy

Cells were plated into a 25 cm^2^ flask (5 × 10^5^ cells/condition). After the indicated time treatment, cells were fixed for 60 min at 4°C in 2% glutaraldehyde in 0.1 M Sörensen phosphate buffer (pH 7.3), post-fixed 1h in aqueous 2% osmium tetroxide, stained in bloc in 2% uranyl acetate in 30% methanol, dehydrated and finally embedded in Epon (Electron Microscopy Sciences, 14120). Ultrathin sections were stained with uranyl acetate and lead citrate and examined with an FEI Tecnai 12 electron microscope. Digital images were taken with a SIS MegaviewIII CCD camera.

### RNA isolation and reverse transcription quantitative PCR

Cells were plated into a 25 cm^2^ flask (2 × 10^6^ cells/condition). After the indicated treatment time, cells were collected by centrifugation. According to the manufacturer’s instructions, their pellets were subjected to RNA extraction using beta-mercaptoethanol and the RNeasy® Plus Mini kit (Qiagen). The genomic DNA was then eliminated using the ezDNase™ enzyme and samples were then subjected to the reverse transcription of the samples (2 µg RNA per sample) using the Master Mix SuperScript™ IV VILO™ kit (Invitrogen, 11766050). A 10-fold dilution of the resulting cDNA was amplified employing SsoAdvanced Univ SYBR Green Supermix (Bio-Rad, 1725272) in a 10 μL volume with the following program: 95°C for 30 seconds, 40 cycles of 95°C for 10 seconds, and 60°C for 20 seconds using the CFX96 Touch Real-Time PCR (Biorad). Primers used for amplification are listed in Table S1 and were purchased from Eurogentec. Quantification of mRNA levels was performed using the ΔΔCt method. Results were normalized to *PPIA* (*Peptidyl Propyl Isomerase A*).

### Tumor growth assessment in immunocompetent C57BL/6 mice

Seven weeks-old female WT C57BL/6 mice were obtained from Envigo (Grannat, France) and were maintained in the animal facilities of the Centre de Recherche des Cordeliers in specific pathogen-free conditions in a temperature-controlled environment with 12 h light/12 h dark cycles. Animal experiments followed the Federation of European Laboratory Animal Science Association guidelines and the EU Directive 63/2010. Protocol APAFIS #36901-2022021116138370v4 was approved by the local Ethical Committee (CEEA C. Darwin n. 5, registered at the French Ministry of Research).

To establish syngeneic solid tumors, 3 × 10^5^ WT MCA205 cells were inoculated subcutaneously (in the back of the mice into immunocompetent WT C57BL/6 female mice, near the thigh. Tumor volume (longest dimension × perpendicular dimension x height dimension) was routinely monitored using a digital caliper. When the tumor volume reached 70 mm3, mice received 5 mg/kg intratumoral (i.t.) CX-3543 (n=9), 3 mg/kg intraperitoneal (i.p.). DC661 (n=9), both 5 mg/kg intratumoral (i.t.) CX-3543 and 3 mg/kg intraperitoneal (i.p.). DC661 (n=11) or an equivalent volume of vehicle (n=11). Mice were all treated with drugs or vehicles at days 7, 8, 9, 13, and 14 post-tumor inoculation. Tumor growth was assessed every two days. Animals bearing necrotic neoplastic lesions or lesions that exceeded 20% to 25% of their body mass were euthanized. Data were analyzed using GraphPad Prism software and https://kroemerlab.shinyapps.io/TumGrowth/.

### Statistical analysis

The exhaustive description of statistical analysis for each figure is provided in a table (Table S4). For FACS and immunofluorescence data analysis, most of the statistical analysis was done within R software [61], version 4.2.1, within RStudio environment [62]. For the analysis of the FACS data, results represent the percentage of dead cells in each replicate and were plotted as median with minimum and maximum values. For immunofluorescence data analysis, results were plotted as the median value with minimum and maximum values. Statistical significance was performed using robust linear models that were fitted with the ‘rlm’ R function of package ‘MASS’ [63]. P-values were then calculated with ‘rob.pvals’ R function of package ‘repmod’. In the case of technical replicates (eg several fields acquired for each well), robust linear mixed models were fitted with ‘rlmer’ R function of package ‘robustlmm’ [64] that provides t-statistics for model coefficients. P-values were then calculated from t-statistics, using Satterthwaite approximation formula for degree of freedom provided by ‘lmer’ R function of package ‘lmerTest’ [65].

For live cell imaging data, statistical significance was calculated as follows:

**Figure 7C**, p-values were obtained from the linear and quadratic coefficient of each curve, applying a 3-coefficient linear model (see Table S4): dots = Inter + Lin * Time + Quad * Time^2^ (coefficients are available in an extended version of **Figure 7C**, Figure S7A). The light green curve’s (corresponding to the CX-3543 fluorescence after blue light excitation) linear coefficient is smaller, and its quadratic coefficient is larger in absolute value than the dark green curve (corresponding to CX-3543 without blue light excitation). Using the CX-3543 without blue light excitation condition as the control, significant p-values indicated differences in these coefficients (p-values < 0.0001). The extremum position, given by (Lin)/(2 * Quad), shows that condition CX-3543 with blue light excitation reaches its maximum significantly earlier than the CX-3543 without blue light excitation condition.

**Figure 7D**, a linear regression was considered: intensity = Inter + Lin * Time (coefficients are available in an extended version of **Figure 7D**, Figure S7B). The slope (Lin coefficient) of the green curve is significantly higher (p-value < 0.0001) than the grey one.

**Figure 7E**, the temporal behavior seems different before 110 min and after. Therefore, we applied two linear regression model, fraction = Inter + Lin * Time, one before 110 min and one after (coefficients are available in an extended version of **Figure 7E**, Figure S7C). Before 110 min, no curves have a negative slope. After 110 min, only the green curves have significant negative slope (p-value < 0.0001 for both); and we observed that the slope of the light green curve is significantly smaller that the dark green one (p-values < 0.0001).

Where indicated, (figures 2c-f, 3i, 4b, 4i-j, 5c-f, 6a-e, S4a, S5f, S6a-c), 2-coefficients linear models with interaction were used. In these cases, p-values associated to the additive effect of both coefficients were used.

For the RT-qPCR data, statistical significance was determined using GraphPad Prism version 10.1.1 software. The two-tailed unpaired Mann-Whitney test was applied to the data obtained from three independent experiments, each performed in triplicates. The results are presented as mean values ± standard deviation (SD).

For the MTT data. Results are expressed as mean values ± standard deviation (SD) of the percentage of cell viability from three wells.

The *in vivo* data were analyzed using the open-access TumGrowth software (https://github.com/kroemerlab/TumGrowth). Results were plotted as mean ± standard error of the mean (SEM). Statistical significance of the survival curve was determined using a log-rank Mantel-Cox test.

## Acknowledgments and funding

The authors thank Sylvie Souquère who conducted electron microscopy studies at the electron microscopy facility AMMICA-UAR3655 of the Institute Gustave Roussy (IGR), Villejuif, France. The authors also thank the staff of CRISP’edit technology platforms (INSERM US 005 – CNRS UAR 3427-TBMCore, Université de Bordeaux, France) for assistance. The authors also thank the staff of the CEF core facility at CRC.

L.F is supported by a doctoral fellowship from the Ecole Doctorale Paris-Saclay. K.A-V is supported by the Mexican National Council of Science and Technology (CONACYT, funding #757821) and by the Ligue Contre le Cancer (IP/SC-17519). M.D-M is supported by funds from the Institut National de la Santé et de la Recherche Médicale (Inserm) and grants from the Agence National de la Recherche (ANR-21-CE44-0016 (CISCO), ANR-23-CE13-0013 (JANUS), INCa (PLBIO23-216-2023-181). J.L.M. and L.G. are funded by Inserm, CNRS, Ecole Polytechnique, and INCa PLBIO (PLBIO 20-117) and ANR G4Access (ANR-20-CE12-0023) grants. GK is supported by the Ligue contre le Cancer (équipe labellisée); Agence National de la Recherche (ANR-22-CE14-0066 VIVORUSH, ANR-23-CE44-0030 COPPERMAC, ANR-23-R4HC-0006 Ener-LIGHT); Association pour la recherche sur le cancer (ARC); Cancéropôle Ile-de-France; Fondation pour la Recherche Médicale (FRM); a donation by Elior; European Joint Programme on Rare Diseases (EJPRD) Wilsonmed; European Research Council Advanced Investigator Award (ERC-2021-ADG, Grant No. 101052444; project acronym: ICD-Cancer, project title: Immunogenic cell death (ICD) in the cancer-immune dialogue); The ERA4 Health Cardinoff Grant Ener-LIGHT; European Union Horizon 2020 research and innovation programmes Oncobiome (grant agreement number: 825410, Project Acronym: ONCOBIOME, Project title: Gut OncoMicrobiome Signatures [GOMS] associated with cancer incidence, prognosis and prediction of treatment response, Prevalung (grant agreement number 101095604, Project Acronym: PREVALUNG EU, project title: Biomarkers affecting the transition from cardiovascular disease to lung cancer: towards stratified interception), Neutrocure (grant agreement number 861878: Project Acronym: Neutrocure; project title: Development of “smart” amplifiers of reactive oxygen species specific to aberrant polymorphonuclear neutrophils for treatment of inflammatory and autoimmune diseases, cancer and myeloablation); National support managed by the Agence Nationale de la Recherche under the France 2030 programme (reference number 21-ESRE-0028, ESR/Equipex+ Onco-Pheno-Screen); Hevolution Network on Senescence in Aging (reference HF-E Einstein Network); Institut National du Cancer (INCa); Institut Universitaire de France; LabEx Immuno-Oncology ANR-18-IDEX-0001; a Cancer Research ASPIRE Award from the Mark Foundation; PAIR-Obésité INCa_1873, the RHUs Immunolife and LUCA-pi (ANR-21-RHUS-0017 and ANR-23-RHUS-0010, both dedicated to France Relance 2030); Seerave Foundation; SIRIC Cancer Research and Personalized Medicine (CARPEM, SIRIC CARPEM INCa-DGOS-Inserm-ITMO Cancer_18006 supported by Institut National du Cancer, Ministère des Solidarités et de la Santé and INSERM). This study contributes to the IdEx Université de Paris Cité ANR-18-IDEX-0001. Views and opinions expressed are those of the author(s) only and do not necessarily reflect those of the European Union, the European Research Council or any other granting authority. Neither the European Union nor any other granting authority can be held responsible for them.

## Abbreviations

ATG7: autophagy related 7
ATG13: autophagy related 13
Baf A1: Bafilomycin A1
CTSB: cathepsin B
DKO: double knockout
KO: knockout
LAMP1: lysosomal associated membrane protein 1
LAMP2: lysosomal associated membrane protein 2
LGALS3: galectin 3
MAP1LC3 (LC3): microtubule associated protein 1 light chain 3 beta
mTORC1: mechanistic target of rapamycin complex 1
POL I: RNA Polymerase I
SQSTM1/p62: sequestosome 1
TFEB: transcription factor EB
TFE3: transcription factor E3
ULK1: unc-51 like autophagy activating kinase 1

