## Supplementary information for "Lysosomal membrane permeabilization enhances the anticancer effects of RNA Polymerase I transcription inhibitors"

### Supplementary Figure 1.

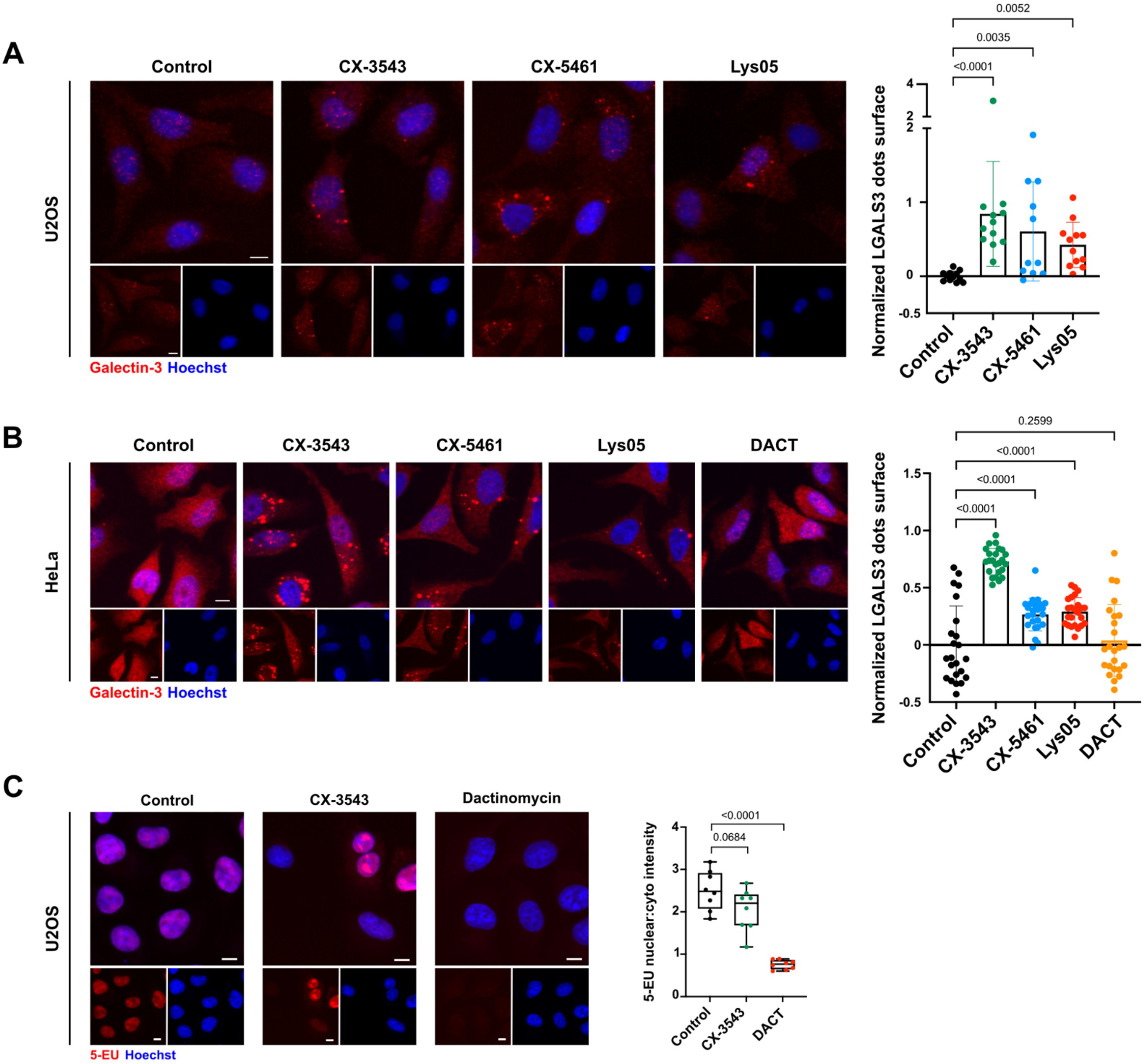


**Figure S1.** CX-3543 and CX-5461 promote lysosomal membrane permeabilization.

**A)** U2OS cells were treated either with CX-3543 (3 μM) or CX-5461 (5 μM) for 16 h and then immunostained for Galectin 3. HCM analysis was then performed to evaluate the endogenous Galectin 3 puncta surface per cell, with a minimum of 2,400 cells analyzed per condition. Lys05 (10 μM, 16 h) was used as a positive control for LMP induction.

**B)** HeLa cells were treated either with CX-3543 (3 μM), CX-5461 (5 μM) for 16 h or Dactinomycin (1 μM, 2 h) and then immunostained for detection of endogenous Galectin 3. HCM was performed to evaluate endogenous Galectin 3 puncta surface per cell, with a minimum of 2,400 cells analyzed per condition. Lys05 (10 μM, 16 h) was used as a positive control for LMP induction.

**C)** U2OS cells were treated either with CX-3543 (3 μM) or Dactinomycin (1 μM, 2 h) and then 5-ethynyl uridine (EU) was added during the last hour of treatment. Dactinomycin (1 μM, 2 h) was used as a positive control for transcription inhibition. HCM analysis was performed to assess the 5-EU nuclear fluorescence intensity, with a minimum of 1,200 cells analyzed per condition. The data are expressed as 5-EU nuclear/cytoplasmic fluorescence intensity ratio which indicates RNA transcription levels.

**Data analysis**: Statistical significance was calculated by means of 1-coefficient robust linear mixed model (S1A-C). Scale bars: 10 μm.

**Supplementary Figure 2.**

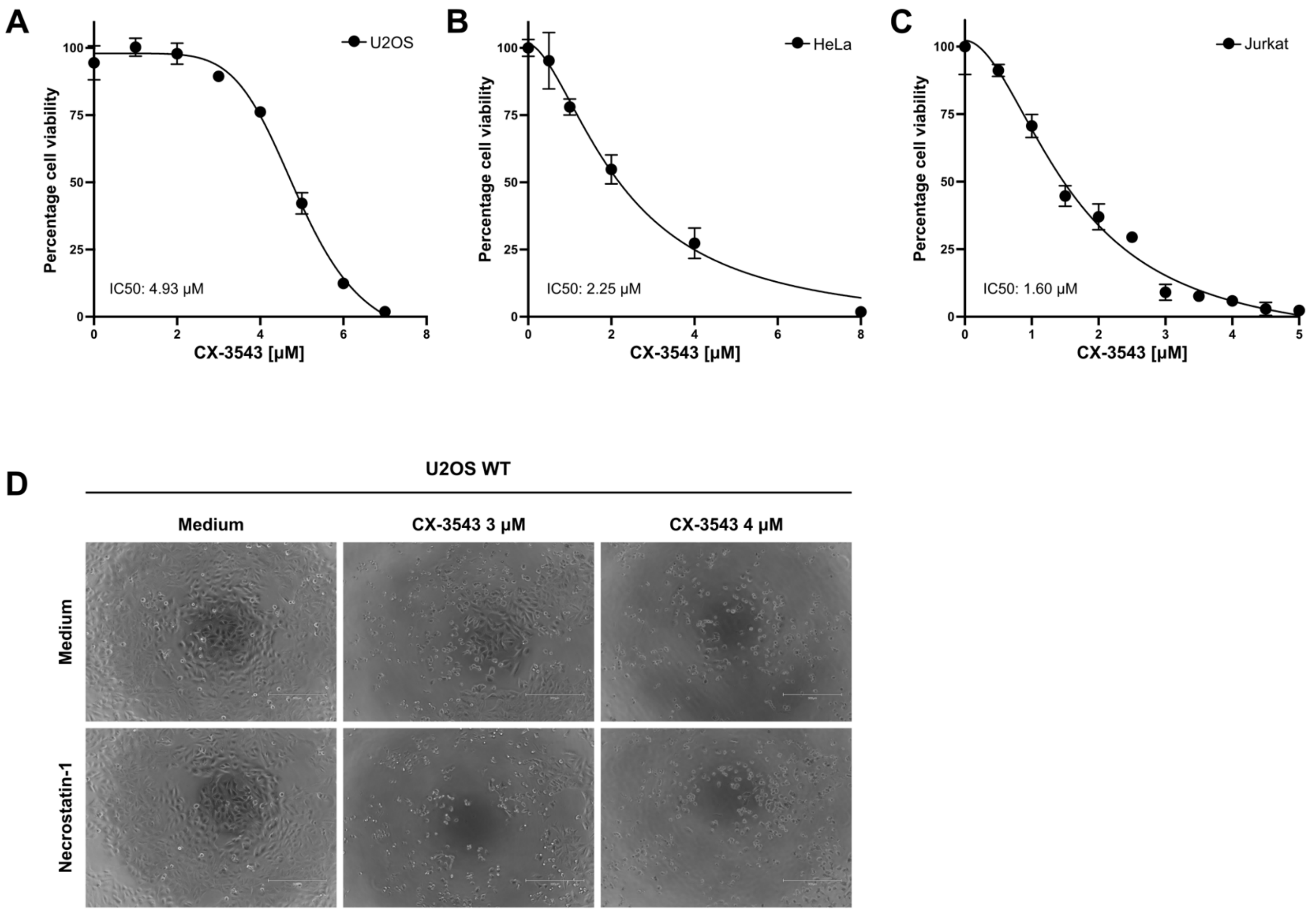


**Figure S2.** CX-3543 reduces cell viability in U2OS, HeLa and Jurkat cancer cell lines.
**A-B)** Cell viability analysis by MTT assay in U2OS (**A**) or in HeLa cells (**B**) after CX-3543 dose escalation treatment (0.5-8 µM) for 24 h.
**C)** Cell viability analysis by MTT assay in Jurkat cells after CX-3543 dose escalation treatment (0.5-5 µM) for 24 h.
**D)** Phase-contrast images from U2OS WT cells that were pre-treated with the necroptosis inhibitor necrostatin-1 (20 µM), for 1 h before introducing 3 µM or 4 µM CX-3543 for 24 h.
**Data analysis**: Statistical significance was calculated by means of a non-linear regression model for S2A, B, C.

### Supplementary Figure 3.

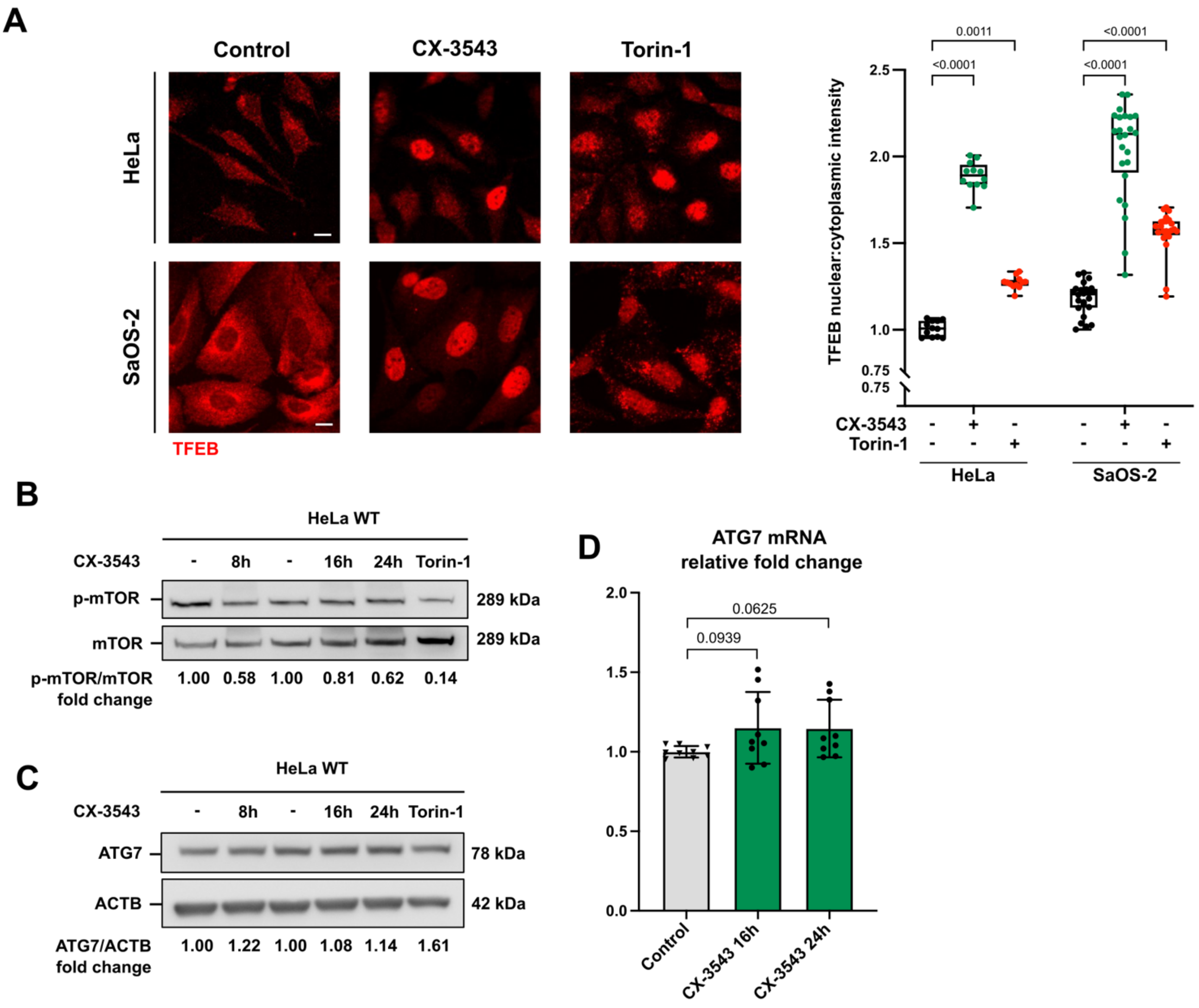


**Figure S3.** Investigating the regulation of TFEB and ATG7 in cells treated with CX-3543.
**A)** HeLa and SaOS-2 cells were treated with 3 µM CX-3543 for 4 h, and then immunostained for detection of endogenous TFEB before being subjected to HCM analysis. Torin-1 (600 nM) was used as a positive control to induce nuclear translocation of TFEB. Left panel: Representative images of immunofluorescence staining of endogenous TFEB in HeLa (upper panel) and SaOS-2 (lower panel) cells. Right panel: Evaluation of the ratio of TFEB nuclear fluorescence intensity to TFEB cytoplasmic fluorescence intensity, with a minimum of 2,400 cells acquired per condition.
**B-C)** Immunoblot analyses of expression levels of mTOR phosphorylation at residue Ser2448 (**B**) and ATG7 (**C**) from HeLa cells treated with 3 µM CX-3543 for indicated time points. Torin-1 (600 nM, 3 h) was used as a potent inhibitor of mTOR. The expression levels of phospho-mTOR and ATG7 were quantified and normalized to mTOR and ACTB levels, respectively. The quantification values represent the ratio of treated cells versus untreated cells (fold change).
**D)** Real-time-quantitative PCR of ATG7 mRNA expression levels in U2OS cells treated with CX-3543 (3 µM) for indicated time points. Data represent values normalized to Peptidyl Propyl Isomerase A (PPIA) housekeeping gene and expressed as fold change (N=3).
**Data analysis**: Statistical significance was calculated by means of 1-coefficient robust linear mixed model (**S3A**) or a two-tailed unpaired Mann-Whitney (**S3D**) model. Scale bars: 10 µm.

### Supplementary Figure 4.

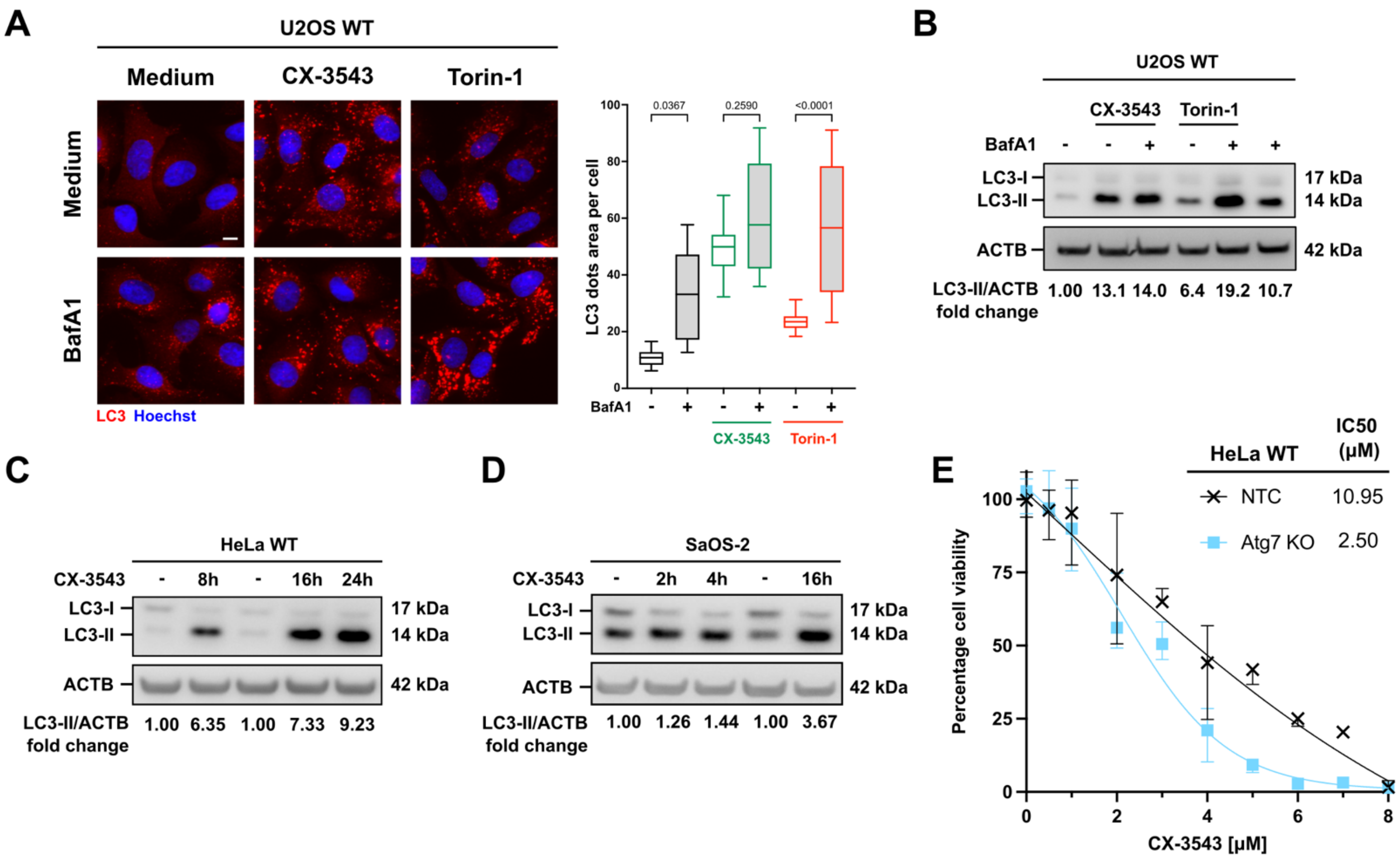


**Figure S4.** Regulation of autophagy by CX-3543.
**A)** U2OS cells were treated with CX-3543 (3 μM) and Bafilomycin A1 (10 nM) for 6 h and then immunostained for detection of endogenous LC3. HCM analysis was performed to evaluate the area of LC3 puncta per cell, with a minimum of 3,200 cells analyzed per condition. Torin-1 (600 nM, 4 h) was used as a potent autophagy inducer.
**B)** Evaluation of the autophagy flux in response to CX-3543. U2OS cells were treated for 2 h with Bafilomycin (50 nM) before adding CX-3543 (3 μM, 6 h). Immunoblot analysis of the expression levels of LC3-II was shown. Band intensities were quantified relative to β-actin and normalized to untreated cells (fold change).
**C-D)** Western blot analyses of the expression of LC3-II levels in HeLa (**C**) and SaOS-2 (**D**) cells treated with CX-3543 (3 μM) for indicated time points. Band intensities were quantified relative to ACTB and normalized to untreated cells (fold change).
**E)** Cell viability measurements in NTC and ATG7 KO HeLa cells after 24 h of treatment with CX-3543 at concentrations 0.5 - 8 μM.
**Data analysis**: Statistical significance was calculated by means of a 2-coefficient robust linear mixed model (**S4A**) or a non-linear regression (**S4E**)” model. Scale bars: 10 μm.

**Supplementary Figure 5.**

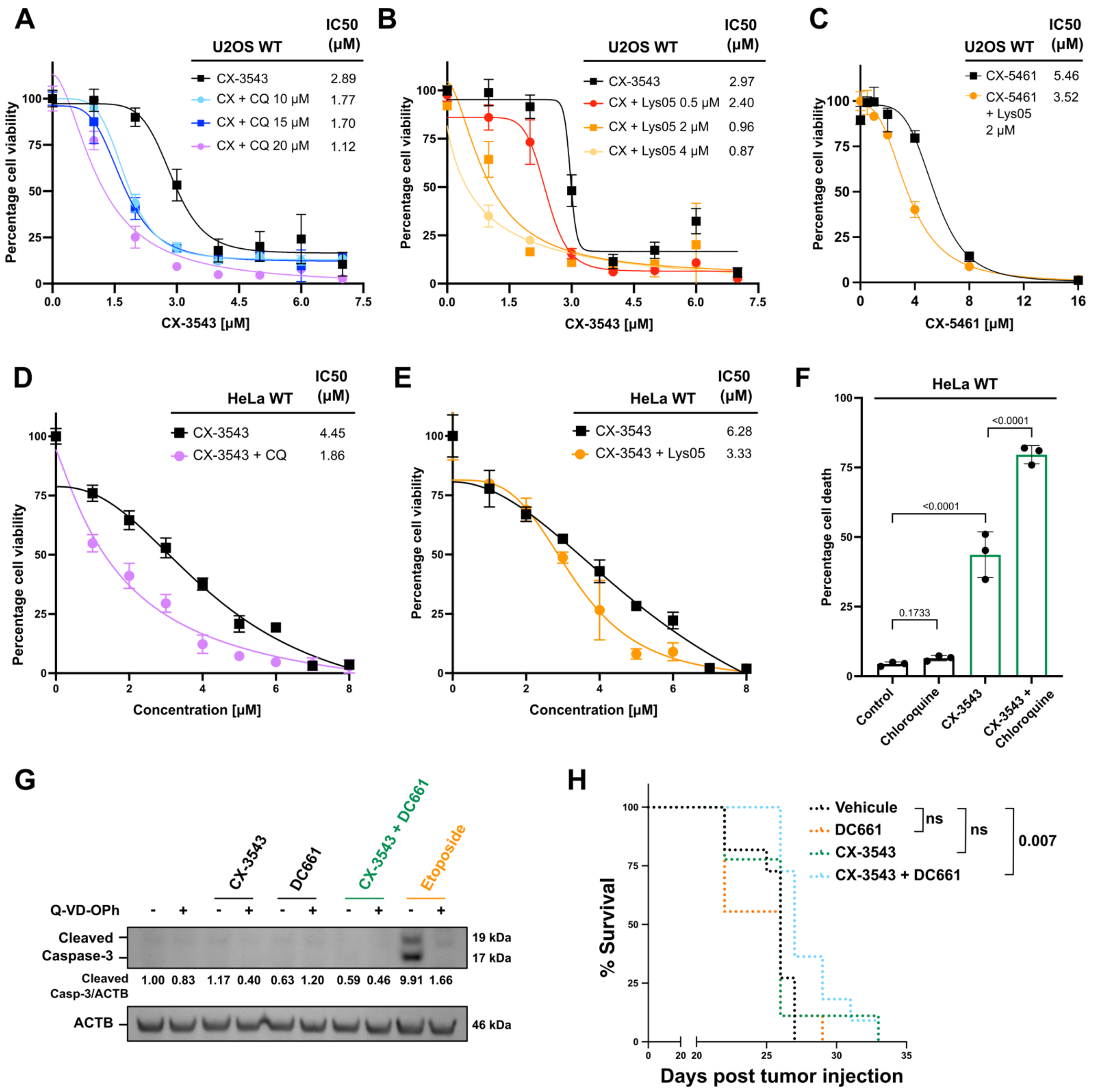


**Figure S5.** Targeting lysosomes enhances the anticancer efficacy of CX-3543 and CX-5461 POL I transcription inhibitors.

**A-B)** Cell viability in U2OS cells was assessed following a 24 h treatment with escalating doses of CX-3543, either alone or in combination with chloroquine or Ly05, at indicated concentrations.

**C)** Cell viability in U2OS cells was assessed following treatment with escalating doses of CX-5461, either alone or in combination with 2 μM Ly05 for 24 h.

**D-E)** Cell viability in HeLa cells was assessed following a 24 h treatment with escalating doses of CX-3543, either alone or in combination with 20 μM chloroquine or 2 μM Ly05.

**F)** Cell death was assessed in HeLa after a 24 h treatment with CX-3543 (2 μM), either alone or in combination with chloroquine (20 μM) using a Live/Dead Far Red assay.

**G)** U2OS cells were pre-treated with Q-VD-OPh (40 μM) 2 h, followed by a 24 h treatment with CX-3543 (2 μM), administrated alone or in combination with DC661 (800 nM). Etoposide (100 μM) was used as an apoptosis inducer. Western blot image of the appearance of the cleaved forms of Caspase-3 was shown. The expression levels of the cleaved Caspase-3 forms were quantified relative to ACTB and normalized to untreated cells.

**H)** Percentage of tumor free mice and overall survival are reported from the in vivo experiment (see Figure 5G).

**Data analysis**: Statistical significance was calculated by means of a non-linear regression model (**S5A-E**), a 2-coefficients robust linear model (**S5F**), or a log-rank Mantel-Cox test (**S5H**).

**Supplementary Figure 6.**

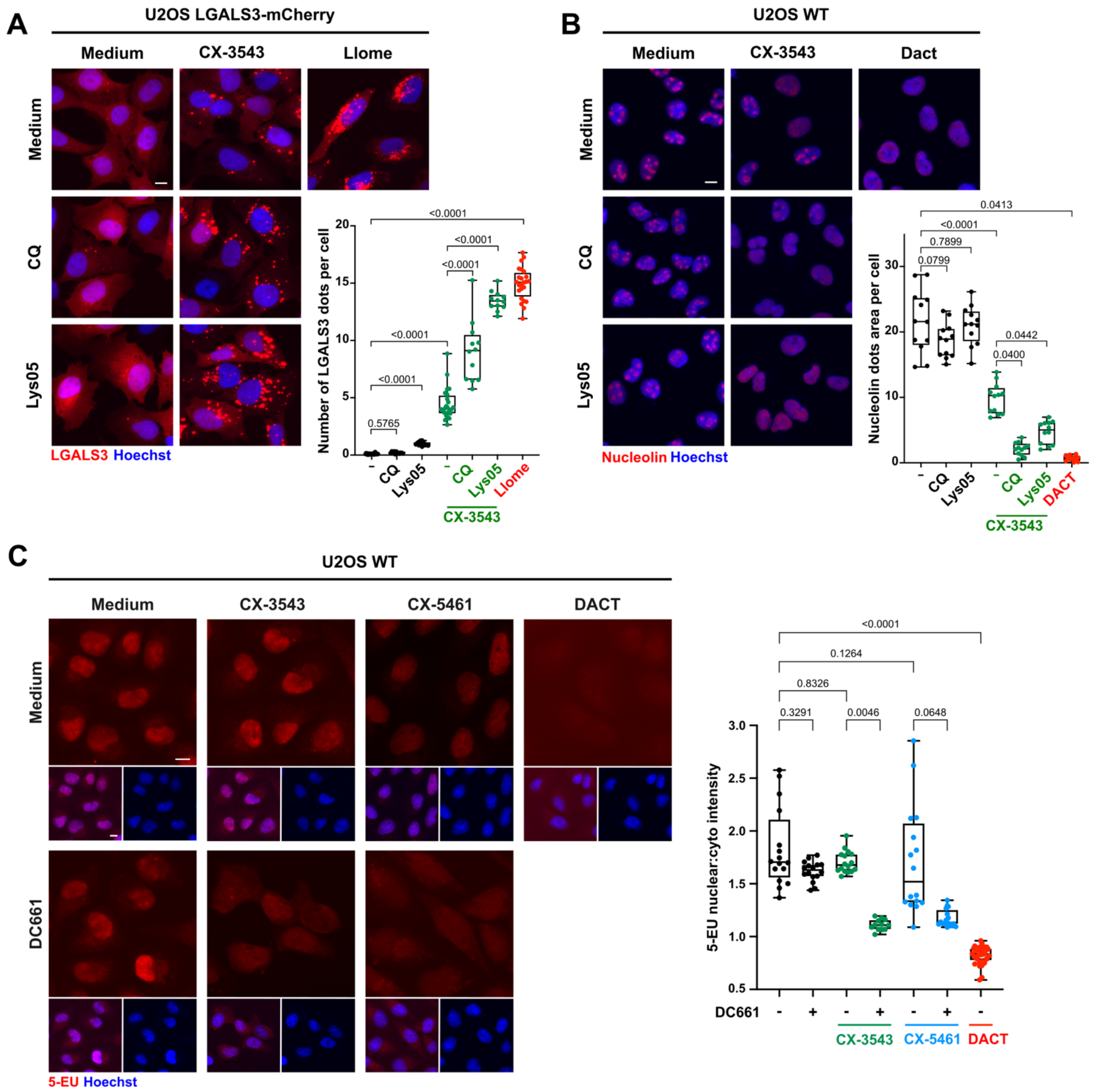


**Figure S6.** Chloroquine derivatives enhance the lysosomal and nuclear toxicity of CX-3543 and CX-5461.

**A)** U2OS cells stably expressing LGALS3-mCherry were treated with CX-3543 (2 µM) either alone or in combination with chloroquine (15 µM) or Lys05 (2 µM). LLOMe (1 mM, 1 h) was used as a positive control for LMP induction. HCM analysis was performed to evaluate the number of Galectin 3 puncta per cell, with a minimum of 2,400 cells analyzed per condition.

**B)** U2OS cells were treated with 2 µM CX-3543 either alone or in combination with chloroquine (15 µM) or Lys05 (2 µM) for 16 h and then immunostained for detection of Nucleolin. HCM analysis was performed to evaluate the area of Nucleolin puncta per cell, with a minimum of 2,400 cells analyzed per condition. Dactinomycin (1 µM, 2 h) was used as an inducer of nucleolar disruption.

**C)** U2OS cells were treated with CX-3543 (2 µM) or CX-5461 (3 µM) either alone or in combination with DC661 (800 nM) for 16 h. 5-ethynyl uridine (EU) was added during the last hour of treatment. Dactinomycin (1 µM, 2 h) was used as a positive control for transcription inhibition. HCM analysis was performed to assess the 5-EU nuclear fluorescence intensity, with a minimum of 1,200 cells analyzed per condition. The data are expressed as 5-EU nuclear/cytoplasmic fluorescence intensity ratio which indicates RNA transcription levels.

**Data analysis**: Statistical significance was determined using a two-coefficients robust linear mixed model followed by a one-coefficient robust linear mixed (**S6A-C**)" model. Scale bars: 10 µm.

### Supplementary Figure 7.

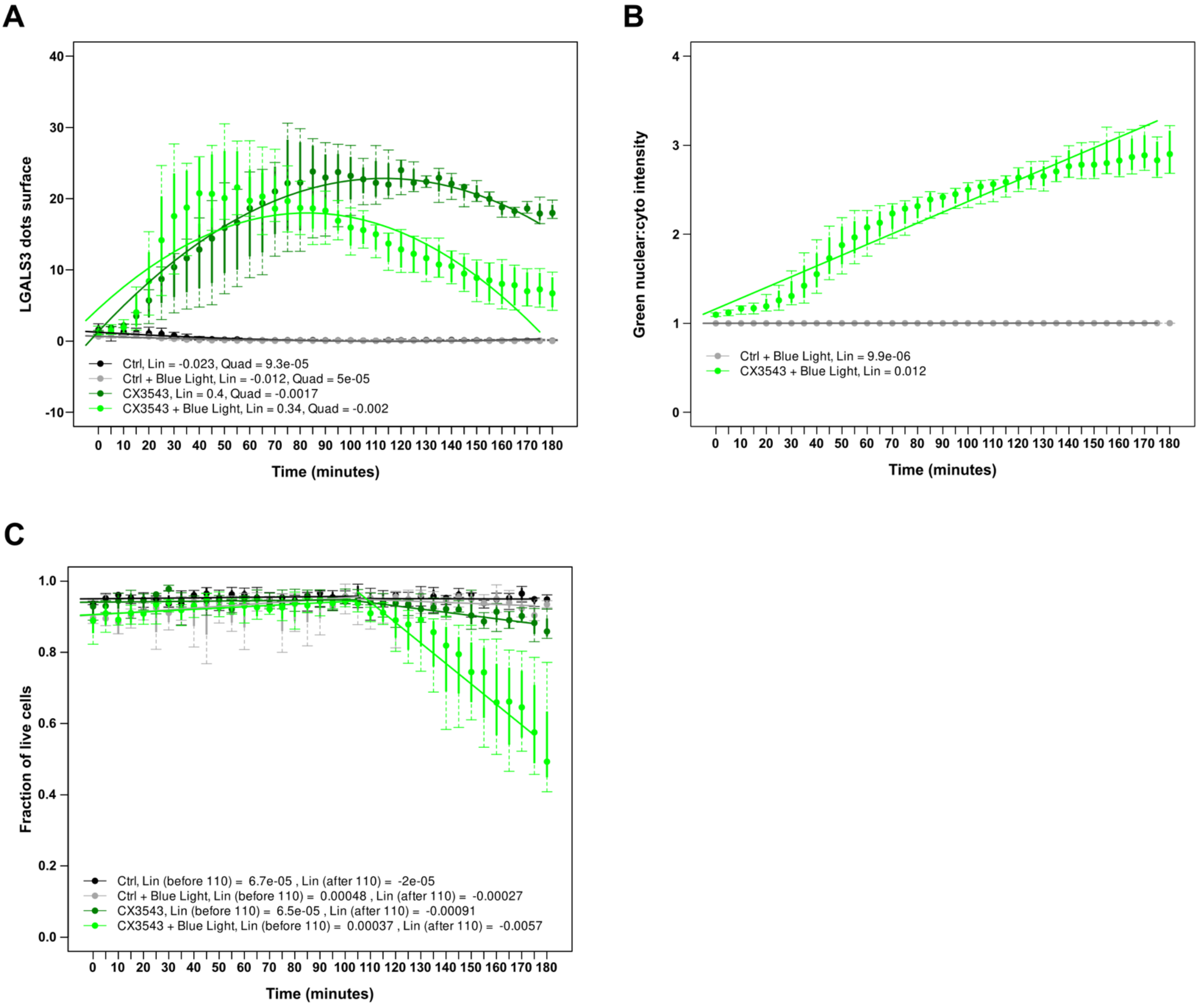


**Figure S7.**
**A)** Extended version of figure 7C with coefficients displayed.
**B)** Extended version of figure 7D with coefficients displayed.
**C)** Extended version of figure 7E with coefficients displayed.
**Data analysis**: Regression analysis and associated coefficients were detailed in Statistical analysis.

**Table S1.** The PCR primer sequences used in the quantitative RT-PCR experiments are shown.

| Target | Forward (5'-3') | Reverse (5'-3') |
|---|---|---|
| *LAMP1* | ACGTTACAGCGTCCAGCTCAT | TCTTTGGAGCTCGCATTGG |
| *LAMP2* | GCTCCTTAACCCAGTGTCATTA | GCTCCTTAACCCAGTGTCATTA |
| *CTSB* | GCAAGGCTCCGTCTCAATAA | TCCCAAAGTGCTGGGATTAC |
| *CTSD* | CATTCCCGAGGTGCTCAAGA | CACGTAGGTGCTGGACTTGT |
| *ATP6V0A1* | GTCCTTCGCCGTCAGTATTT | GTCCTTCGCCGTCAGTATTT |
| *ATP6V0D1* | TGTGCGTGTGTGTGTATGT | GTCTCTAAGAGGGTCAGGAGAA |
| *ATP6V0E1* | CATTGTGATGAGCGTGTTCTGG | AACTCCCCGGTTAGGACCCTTA |
| *MAP1LC3B* | AAGGCGCTTACAGCTCAATG | CTGGGAGGCATAGACCATGT |
| *SQSTM1* | GCCTCTGGTTCTGACACTTT | GGTGAGGTGGAAGGCATTTA |
| *ATG7* | GCGTACGAGGAAGCGTATAAA | GTGTCTTCCAACTTCCCTACAT |
| *PPIA* | CACCGCCGAGGAAAACCG | AAAATTTTCTGCTGTCTTTGGGA |

**Table S2.** The isobologram table summarizes the various drug concentrations (both alone and in combination) and the corresponding Combination Indexes (CI) for a given lethal dose. For each increasing Lethal Dose (LD), the table indicates the concentrations required to achieve the effect, whether through single or combinational treatment. The Combination index (CI) derived from these values is listed in the final column.

| Lethal dose (LD) | [CX-3543] | [DC661] | [CX-3543 in mix] | [DC661 in mix] | CI |
|---|---|---|---|---|---|
| 0.6 | 5.373458 | 1.505875 | 1.68884 | 0.675536 | 0.762893 |
| 0.5 | 5.236768 | 1.315324 | 1.556816 | 0.622726 | 0.770725 |
| 0.4 | 5.109697 | 1.217257 | 1.428503 | 0.571401 | 0.748984 |
| 0.3 | 4.974983 | 1.13024 | 1.282498 | 0.512999 | 0.711675 |
| 0.2 | 4.805695 | 1.014796 | 1.053128 | 0.421251 | 0.634251 |

**Table S3.** The sequences of single guide (sg) RNA that specifically target TFEB or TFE3 used for the lentiviral vector production are shown. The PCR primers used for amplifying the DNA region of interest prior to Sanger sequencing are listed in the table.

| Gene | gRNA name | Target sequence (5'-3') | PCR primers couples (5'-3') |
|---|---|---|---|
| *TFEB* | TFEB1-G1 | GAAGTGGACGGGGGTATTGA | CTGCCCCTTCTCATCCACAG / GGGCTATTGGGAGCACTGTT |
| *TFE3* | TFE3-G2 | GGGGTGGACGGCTCAATGTG | CCATTCCACAGCCTCCCAAT/ CTCATGAGCCTCCATGGTCC |

**Table S4.** Detailed statistical tests used in this study.

| Figures | Software | Variables | StatisticalModels | Rfunctions |
|---|---|---|---|---|
| 1b | GraphPad Prism V10.1.1 | green, red, blue channels | Pearson R Correlation Factor | N.A |
| 1c | GraphPad Prism V10.1.1 | green, red, blue channels | Pearson R Correlation Factor | N.A |
| 1f | R software | condition, well | 1-coefficient robust linear mixed model with Satterthwaite approximation formula | rlmer(value ~ condition + (1\|well)) |
| 1h | R software | condition, well | 1-coefficient robust linear mixed model with Satterthwaite approximation formula | rlmer(value ~ condition + (1\|well)) |
| 1i | R software | condition, well | 1-coefficient robust linear mixed model with Satterthwaite approximation formula | rlmer(value ~ condition + (1\|well)) |
| 2b | R software | condition | 1-coefficient robust linear model with Satterthwaite approximation formula | rlm(value ~ condition) |
| 2c | R software | treatment, condition, well | 2-coefficient robust linear mixed model with Satterthwaite approximation formula | rlmer(value ~ treatment / condition + (1\|well))) |
| 2d | R software | treatment, condition, well | 2-coefficient robust linear mixed model with Satterthwaite approximation formula | rlmer(value ~ treatment / condition + (1\|well))) |
| 2e | R software | treatment, condition, well | 2-coefficient robust linear mixed model with Satterthwaite approximation formula | rlmer(value ~ treatment / condition + (1\|well))) |
| 2f | R software | treatment, condition, well | 2-coefficient robust linear mixed model with Satterthwaite approximation formula | rlmer(value ~ condition / treatment + (1\|well))) |
| 3b | R software | condition, well | 1-coefficient robust linear mixed model with Satterthwaite approximation formula | rlmer(value ~ condition + (1\|well)) |
| 3b | R software | condition, well | 1-coefficient robust linear mixed model with Satterthwaite approximation formula | rlmer(value ~ condition + (1\|well)) |
| 3d | GraphPad Prism V10.1.1 | treatment | two-tailed unpaired Mann-Whitney test | N.A |
| 3f | R software | condition, well | 1-coefficient robust linear mixed model with Satterthwaite approximation formula | rlmer(value ~ condition + (1\|well)) |
| 3h | GraphPad Prism V10.1.1 | treatment | Non linear regression analysis | N.A |
| 3i | R software | condition, cellLine, well | 2-coefficient robust linear mixed model with Satterthwaite approximation formula | rlmer(value ~ condition / cellLine+ (1\|well))) |
| 4b | R software | treatment, condition, well | 2-coefficient robust linear mixed model with Satterthwaite approximation formula | rlmer(value ~ condition / treatment + (1\|well))) |
| 4b | R software | treatment, condition, well | 2-coefficient robust linear mixed model with Satterthwaite approximation formula, Torin as control treatment | rlmer(value ~ condition / treatment + (1\|well))) |
| 4g | R software | condition, well | 1-coefficient robust linear mixed model with Satterthwaite approximation formula | rlmer(value ~ condition + (1\|well)) |
| 4h | R software | condition, well | 1-coefficient robust linear mixed model with Satterthwaite approximation formula | rlmer(value ~ condition + (1\|well)) |
| 4i | R software | condition, cellLine, well | 2-coefficient robust linear mixed model with Satterthwaite approximation formula | rlmer(value ~ condition / cellLine+ (1\|well))) |
| 4j | R software | condition, cellLine, well | 2-coefficient robust linear mixed model with Satterthwaite approximation formula | rlmer(value ~ condition / cellLine+ (1\|well))) |
| 5c | R software | treatment, condition, well | 2-coefficient robust linear mixed model with Satterthwaite approximation formula | rlmer(value ~ condition / treatment + (1\|well))) |
| 5d | R software | treatment, condition, well | 2-coefficient robust linear mixed model with Satterthwaite approximation formula | rlmer(value ~ condition / treatment + (1\|well))) |
| 5e | R software | treatment, condition, well | 2-coefficient robust linear mixed model with Satterthwaite approximation formula | rlmer(value ~ condition / treatment + (1\|well))) |
| 5f | R software | treatment, condition, well | 2-coefficient robust linear mixed model with Satterthwaite approximation formula | rlmer(value ~ condition / treatment + (1\|well))) |
| 5h | TumGrowth (https://github.com/kroemerlab/TumGrowth) | time, treatment | longitudinal analysis (type II ANOVA and pairwise comparisons across groups) | N.A |
| 6a | R software | treatment, condition, well | 2-coefficient robust linear mixed model with Satterthwaite approximation formula | rlmer(value ~ condition * treatment + (1\|well)) |
| 6a | R software | treatment, well | 1-coefficient robust linear mixed model with Satterthwaite approximation formula, for LLOMe effect | rlmer(value ~ treatment + (1\|well)) |
| 6b | R software | treatment, condition, well | 2-coefficient robust linear mixed model with Satterthwaite approximation formula | rlmer(value ~ condition * treatment + (1\|well)) |
| 6c | R software | time, condition, well | 2-coefficient robust linear mixed model with Satterthwaite approximation formula | rlmer(value ~ (time/condition) + (1\|well)) |
| 6d | R software | treatment, condition, well | 2-coefficient robust linear mixed model with Satterthwaite approximation formula | rlmer(value ~ condition * treatment + (1\|well)) |
| 6d | R software | treatment, well | 1-coefficient robust linear mixed model with Satterthwaite approximation formula, for DACT effect | rlmer(value ~ treatment + (1\|well)) |
| 6e | R software | treatment, condition, well | 2-coefficient robust linear mixed model with Satterthwaite approximation formula | rlmer(value ~ condition * treatment + (1\|well)) |
| 6e | R software | treatment, well | 1-coefficient robust linear mixed model with Satterthwaite approximation formula, for DACT effect | rlmer(value ~ treatment + (1\|well)) |
| 7c | R software | time, quatratic_time, condition | 3-coefficient robust linear model | rlm(value ~ condition / (time + quadratic_time)), rob.pvals |
| 7c | R software | time, quatratic_time, condition | 3-coefficient robust linear model, CX-3543 as control condition | rlm(value ~ condition * (time + quadratic_time)), rob.pvals |
| 7d | R software | time, condition | 2-coefficients robust linear model | rlm(value ~ condition * (time)), rob.pvals |
| 7e | R software | time, condition | 2-coefficients robust linear model, before and after 110 minutes. | rlm(value ~ condition / (time)), rob.pvals |
| 7e | R software | time, condition | 2-coefficients robust linear model, after 110 minutes, CX-3543 as control condition | rlm(value ~ condition * (time)), rob.pvals |

**Table S4.** Detailed statistical tests used in this study.

| Figures | Software | Variables | StatisticalModels | Rfunctions |
|---|---|---|---|---|
| S1a | R software | condition, well | 1-coefficient robust linear mixed model with Satterthwaite approximation formula | rlmer(value ~ (condition) + (1\|well)) |
| S1b | R software | condition, well | 1-coefficient robust linear mixed model with Satterthwaite approximation formula | rlmer(value ~ condition + (1\|well)) |
| S1c | R software | condition, well | 1-coefficient robust linear mixed model with Satterthwaite approximation formula | rlmer(value ~ condition + (1\|well)) |
| S2a | GraphPad Prism V10.1.1 | treatment | Non linear regression analysis | N.A |
| S2b | GraphPad Prism V10.1.1 | treatment | Non linear regression analysis | N.A |
| S2c | GraphPad Prism V10.1.1 | treatment | Non linear regression analysis | N.A |
| S3a | R software | condition, well | 1-coefficient robust linear mixed model with Satterthwaite approximation formula | rlmer(value ~ condition + (1\|well)) |
| S3d | GraphPad Prism V10.1.1 | treatment | two-tailed unpaired Mann-Whitney test | N.A |
| S4a | R software | treatment, condition, well | 2-coefficient robust linear mixed model with Satterthwaite approximation formula | rlmer(value ~ (treatment/condition) + (1\|well)) |
| S4e | GraphPad Prism V10.1.1 | treatment | Non linear regression analysis | N.A |
| S5a | GraphPad Prism V10.1.1 | treatment | Non linear regression analysis | N.A |
| S5b | GraphPad Prism V10.1.1 | treatment | Non linear regression analysis | N.A |
| S5c | GraphPad Prism V10.1.1 | treatment | Non linear regression analysis | N.A |
| S5d | GraphPad Prism V10.1.1 | treatment | Non linear regression analysis | N.A |
| S5e | GraphPad Prism V10.1.1 | treatment | Non linear regression analysis | N.A |
| S5f | R software | treatment, condition, well | 2-coefficients robust linear model | rlm(value ~ condition * treatment), rob.pvals |
| S5h | TumGrowth (https://github.com/kroemer lab/TumGrowth) | time, treatment | log-rank Mantel-Cox test (survival analysis) | N.A |
| S6a | R software | treatment, condition, well | 2-coefficient robust linear mixed model with Satterthwaite approximation formula | rlmer(value ~ condition * treatment + (1\|well)) |
| S6a | R software | treatment, well | 1-coefficient robust linear mixed model with Satterthwaite approximation formula, for LLOMe effect | rlmer(value ~ treatment + (1\|well)) |
| S6b | R software | treatment, condition, well | 2-coefficient robust linear mixed model with Satterthwaite approximation formula | rlmer(value ~ condition * treatment + (1\|well)) |
| S6b | R software | treatment, well | 1-coefficient robust linear mixed model with Satterthwaite approximation formula, for DACT effect | rlmer(value ~ treatment + (1\|well)) |
| S6c | R software | treatment, condition, well | 2-coefficient robust linear mixed model with Satterthwaite approximation formula | rlmer(value ~ condition * treatment + (1\|well)) |
| S6c | R software | treatment, well | 1-coefficient robust linear mixed model with Satterthwaite approximation formula, for DACT effect | rlmer(value ~ treatment + (1\|well)) |
| S7a | R software | time, quatratic_time, condition | 3-coefficient robust linear model | rlm(value ~ condition / (time + quadratic_time)), rob.pvals |
| S7a | R software | time, quatratic_time, condition | 3-coefficient robust linear model, CX-3543 as control condition | rlm(value ~ condition * (time + quadratic_time)), rob.pvals |
| S7b | R software | time, condition | 2-coefficients robust linear model | rlm(value ~ condition * (time)), rob.pvals |
| S7c | R software | time, condition | 2-coefficients robust linear model, before and after 110 minutes. | rlm(value ~ condition / (time)), rob.pvals |
| S7c | R software | time, condition | 2-coefficients robust linear model, after 110 minutes, CX-3543 as control condition | rlm(value ~ condition * (time)), rob.pvals |
